# Host-mediated, cross-generational intraspecific competition in a herbivore species

**DOI:** 10.1101/2020.07.30.228544

**Authors:** Bastien Castagneyrol, Inge van Halder, Yasmine Kadiri, Laura Schillé, Hervé Jactel

## Abstract

Conspecific insect herbivores co-occurring on the same host plant interact both directly through interference competition and indirectly through exploitative competition, plant-mediated interactions and enemy-mediated interactions. However, the situation is less clear when the interactions between conspecific insect herbivores are separated in time within the same growing season, as it is the case for multivoltine species. We hypothesized that early season herbivory would result in reduced egg laying and reduced performance of the next generation of herbivores on previously attacked plants. We tested this hypothesis in a choice experiment with box tree moth females (*Cydalima perspectalis* Walker, Lepidoptera: Crambidae). These females were exposed to box trees (*Buxus sempervirens* L., Buxaceae) that were either undamaged or attacked by conspecific larvae earlier in the season. We then compared the performance of the next generation larvae on previously damaged *vs* undamaged plants. Previous herbivory had no effect on oviposition behaviour, but the weight of next generation larvae was significantly lower in previously damaged plants. There was a negative correlation between the number of egg clutches laid on plants by the first generation and the performance of the next generation larvae. Overall, our findings reveal that early season herbivory reduces the performance of conspecific individuals on the same host plant later in the growing season, and that this time-lagged intraspecific competition results from a mismatch between the oviposition preference of females and the performance of its offspring.

## Introduction

Biotic interactions are strong factors affecting the fitness of interacting individuals, even when interactions are delayed in time or do not imply direct contact between individuals. Such interactions can be found in both plants through plant-soil feedbacks (Putten et al., 2016) and in animals (Fisher et al., 2019; Pfennig & Pfennig, 2020). For instance, insect herbivores exploiting the same plant can compete for food, even when interactions among individuals are separated in time (Kaplan & Denno, 2007). Insects may reduce the impact of interspecific competition by avoiding crowded plants, or plants that have been previously consumed by herbivores, which assumes that they can detect competitors or their effects on plants (De Moraes et al., 2001; Shiojiri & Takabayashi, 2003). For many species, the choice of the oviposition site by mated females is crucial in this respect. The preference-performance hypothesis — *aka* the ‘*mother knows best hypothesis*’ — states that female insects evolved host searching behaviour that leads them to oviposit on hosts where their offspring do best (Gripenberg et al., 2010). A good match between the preference of a mated female for a given plant and the performance of its offspring developing on the same plant implies that females can recognize cues that correlate with larval performance, for instance those related to plant defenses and nutritional quality. Yet, these cues can be largely modified by the simultaneous or sequential presence of other competing herbivores (Bultman & Faeth, 1986; Nykänen & Koricheva, 2004; Abdala‐ Roberts et al., 2019; Visakorpi et al., 2019). Therefore, initialRoberts et al., 2019; Visakorpi et al., 2019). Therefore, initial herbivory by a given species may have time-lagged consequences on the preference and performance of herbivores of another species that subsequently attack the same plant in the same growing season (Poelman et al., 2008; Stam et al., 2014). However, while such time-lagged *interspecific* interactions between herbivores have long been documented (Faeth, 1986), surprisingly much less is known about delayed *intraspecific* interactions in multivoltine species having several generations per year.

Previous herbivory generally reduces the performance of later arriving herbivores on the same plant through different processes. First, the initial consumption of plant biomass can deplete the resource available to forthcoming herbivores, therefore leading to exploitative competition between first and subsequent herbivores (Kaplan & Denno, 2007). Second, initial herbivory triggers a hormonal response that results in the induction and production of anti-herbivore defenses as well as in resource reallocation in plant tissues (Marchand & McNeil, 2004; Blenn et al., 2012; Fatouros et al., 2012; Hilker & Fatouros, 2015; Abdala‐ Roberts et al., 2019; Visakorpi et al., 2019). Therefore, initialRoberts et al., 2019), which generally reduces plant quality and thereby the performance of late coming herbivores (Wratten et al., 1988; Agrawal, 1999; Abdala‐ Roberts et al., 2019; Visakorpi et al., 2019). Therefore, initialRoberts et al., 2019). Such an effect has long been documented in interspecific interactions (Kaplan & Denno, 2007; Moreira et al., 2018), but also in intraspecific interactions. For instance, prior damage by the western tent caterpillar *Malacosoma californicum* Packard (Lepidoptera: Lasiocampidae) induces the regrowth of tougher leaves acting as physical defenses and reducing the fitness of the next tent caterpillars generation (Barnes & Murphy, 2018). Although less common, the opposite phenomenon whereby initial herbivory facilitates damage by subsequent herbivores has also been reported (Sarmento et al., 2011; Godinho et al., 2016; Moreira et al., 2018).

Previous herbivory can also affect the oviposition preference of herbivores that arrive later. Several studies have demonstrated that mated females can discriminate between host plants that have been previously attacked by insect herbivores (Wise & Weinberg, 2002; Stam et al., 2014; Moura et al., 2017; Barnes & Murphy, 2018; Moreira et al., 2018; Weeraddana & Evenden, 2019), thereby reducing competition between herbivores separated in time. Mated females can directly detect the present, past and possibly future presence of competitors themselves. For instance, Averill & Prokopy (1987) showed that female *Rhagoletis pomonella* Walsh (Diptera: Tephritidae) marks its oviposition site with an epideictic pheromone that deters conspecific females from laying eggs, thus reducing intraspecific competition at the larval stage. The frass of several Lepidoptera species was also found to act as an oviposition deterrent (Jones & Finch, 1987; Hashem et al., 2013; Molnár et al., 2017). Mated females may also detect herbivory-induced changes in the physical and chemical characteristics of attacked plants, and consequently avoid laying eggs on less suitable plants. However, several authors reported a mismatch between prior herbivory effects on female oviposition preference *vs* larval growth, consumption or survival of their offspring (Wise & Weinberg, 2002; Bergamini & Almeida-Neto, 2015; Martinez et al., 2017; Godinho et al., 2020). For instance, Weeraddana and Evenden (2019) found that herbivory by the diamondback moth, *Plutella xylostella* (L.) (Lepidoptera: Plutellidae) on canola plants (*Brassica napus* L.) had no effect on subsequent oviposition by the bertha armyworm, *Mamestra configurata* Walker (Lepidoptera: Noctuidae) whereas its larvae had reduced growth on previously damaged plants. Thus, in order to quantify the effect of prior herbivory on subsequent herbivore performance, we need to assess how it affects both female choice and progeny performance in attacked and non-attacked hosts.

In the present study, we investigated the consequences of box tree (*Buxus* spp.) defoliation by the first generation of the box tree moth (BTM) *Cydalima perspectalis* Walker (Lepidoptera: Crambidae) larvae on (*i*) the oviposition behaviour of the adults emerging from those larvae and (*ii*) on the larval performance in the next generation. Specifically, we hypothesized that plants that had previously been attacked by conspecific larvae would (i) receive fewer eggs (*i*.*e*. reduced preference) and (ii) host smaller larvae and chrysalis (*i*.*e*. reduced performance) of the next generation than previously undamaged plants. Our experimental design allowed us to separate the effects of previous herbivory on both preference and performance of conspecific herbivores attacking the same plant in sequence. By doing so, our study brings new insights into the understanding of cross-generational intraspecific competition in insect herbivores and further challenges the ‘*mother knows best hypothesis*.’

## Methods

### Natural history

The BTM is a multivoltine moth species introduced to Europe in 2007 from Asia (Wan et al., 2014). In its native range, BTM larvae can feed on different host genera, whereas in Europe they feed exclusively on box trees (Wan et al., 2014). In the introduced area, BTM larvae overwinter in cocoons tied between two adjacent leaves, mainly in the third instar. Therefore, defoliation restarts in early spring at the beginning of the growing season. In Europe, damage is aggravated by the fact that the BTM has 3-4 generations a year (Kenis et al., 2013; Matošević et al., 2017). When several pest generations successively defoliate the same box tree, there are no leaves left to eat and the caterpillars then feed on the bark, which can lead to the death of the host tree (Kenis et al., 2013; Wan et al., 2014; Alkan Akinci & Kurdoğlu, 2019).

### Biological material

In spring 2019, we obtained box trees from a commercial nursery and kept them in a greenhouse at INRAE Bordeaux forest research station. Box trees were on average 25 cm high and 20 cm wide. We transferred them into 5 L pots with horticultural loam. For two months, we watered them every four days from the above (*i*.*e*. watering leaves too) to remove any potential pesticide remain.

We initiated BTM larvae rearing with caterpillars collected in the wild in early spring 2019, corresponding to those that had overwintered. No ethical approval for animal experimentation was needed. We reared larvae at room temperature in 4320 cm^3^ plastic boxes, and fed them *ad libitum*, with branches collected on box trees around the laboratory. We used the next generation larvae to induce herbivory on box tree plants (experimental treatment, see below) and the subsequent adults for the oviposition experiment. At 25°C, the larval phase lasts for about 30 days and the BTM achieves one generation in 45 days. Adults live 12-15 days. A single female lays on average 800 eggs.

### Experimental design

On June 18^th^ 2019, we haphazardly assigned box trees to *control* and *herbivory* experimental groups. The *herbivory* treatment consisted of *n* = 60 box trees that received five L3 larvae each. Larvae were allowed to feed freely for one week, after which we removed them all from plants. In order to confirm that the addition of BTM larvae caused herbivory, we visually estimated BTM herbivory as the percentage of leaves consumed by BTM larvae per branch, looking at every branch on every plant. We then averaged herbivory at the plant level. Herbivory data were missing in 8 plants. We removed these plants from the analysis testing the effect of prior herbivory as a continuous variable on BTM preference and performance. In the herbivory treatment, the percentage of leaves consumed by BTM larvae ranged from 2.2 to 17.2% and was on average 9.1%. The *control* group (*n* = 61) did not receive any BTM larva. On July 8^th^, we randomly distributed plants of the *herbivory* and *control* treatments on a 11 × 11 grid in a greenhouse (*i*.*e*. total of 121 plants). We left 40 cm between adjacent pots, which was enough to avoid any physical contact between neighbouring plants **(Figure 1, Figure 2)**.

**Figure 1:**
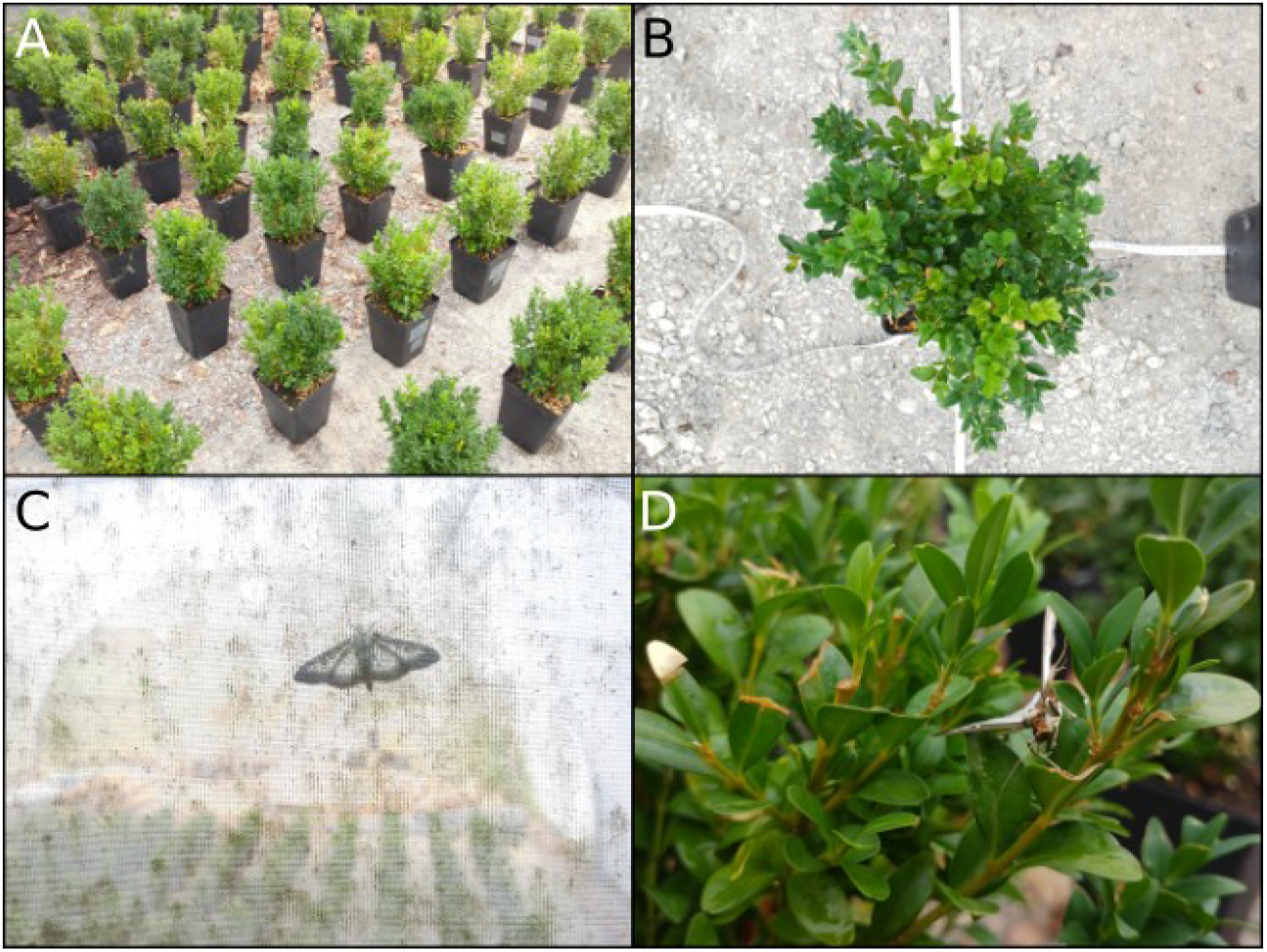
The study design and model species. The two top photos (A, B) illustrate the experimental design and in particular distance among potted plants. Photo C is a view of the greenhouse from the outside, with an adult box tree moth in the foreground, and potted plants in the background. Photo D shows an adult box tree moth on a box tree branch, shortly after it was released.

**Figure 2:**
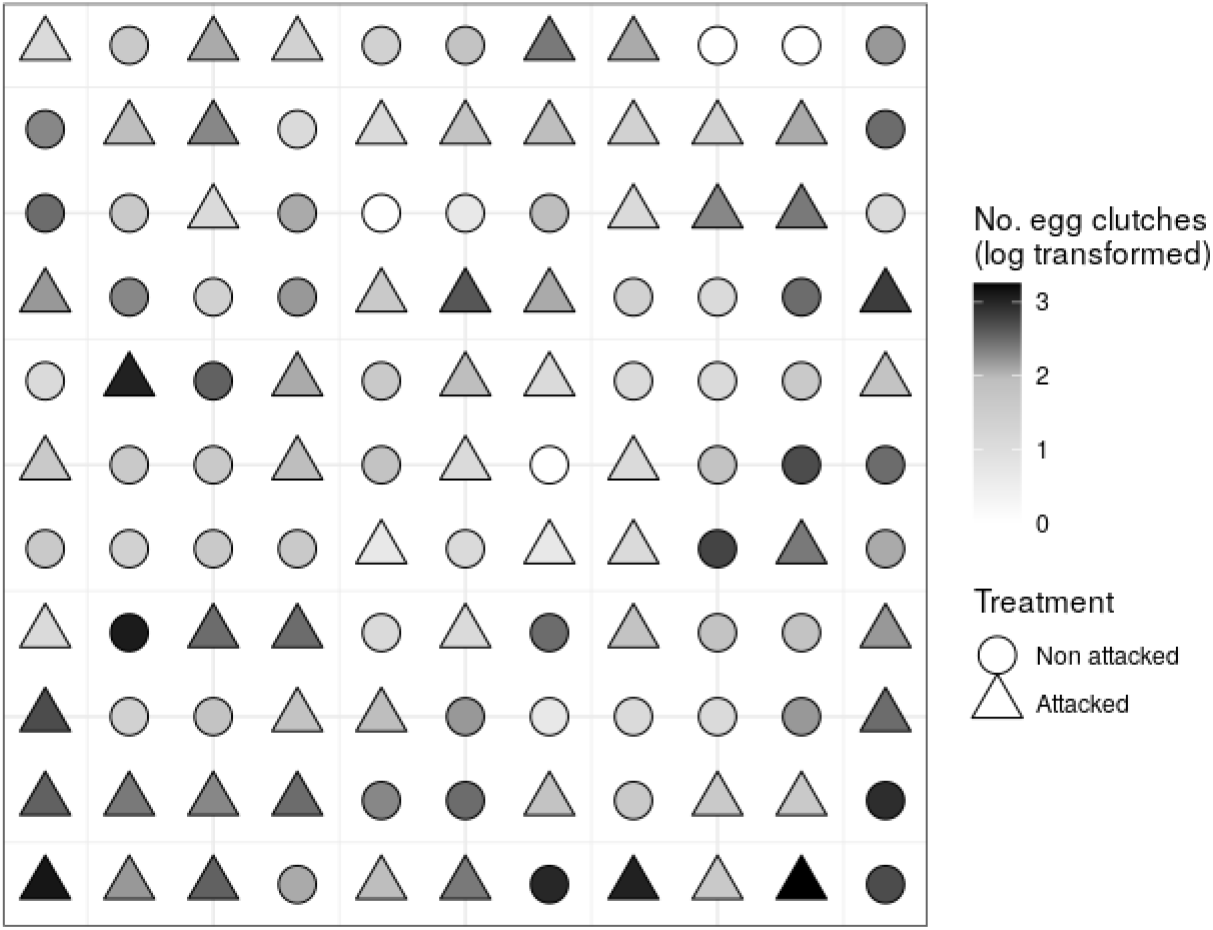
Experimental design. Pots were 40 cm apart. Circles and triangles represent non-attacked (control) and attacked trees. Scale colour represents the number of egg clutches per box tree (log-transformed).

The same day, we released *ca* 100 BTM moths that had emerged from chrysalis less than two days before (*i*.*e*., an uncontrolled mix of males and females). We released moths at the four corners of the experiment to reduce the risk of spatial aggregation. Moths were allowed to fly freely within the greenhouse. They could feed on small pieces of cotton imbibed with a sugar-water solution, disposed on the ground in the greenhouse.

It is important to note that at the time we released moths, there were no larvae feeding on experimental box trees anymore. In addition, at this time, plants in the herbivory treatment had been cleared of caterpillars for three weeks (corresponding to the duration of the chrysalis stage) during which they were watered every two to three days from above. Although larval frass may have been present in pots submitted to the herbivory treatment, it should have been washed out from leaves. Finally, we carried out our experiment in an enclosed greenhouse in which the potential effect of natural enemies on BTM behaviour can be neglected. The consequences are that any effect of prior herbivory on subsequent oviposition behaviour and larval performance should have been independent of cues emitted by BTM larvae themselves or by their frass (Sato et al., 1999; Molnár et al., 2017) and therefore were only plant-mediated.

### BTM host choice

In order to test whether initial defoliation of focal plants influenced host choice for oviposition by BTM females, we counted egg clutches on every branch of every box tree on July 17^th^. Once eggs were counted, we moved box trees to another greenhouse. To prevent larvae from moving from one potted plant to another, we installed box trees in plastic saucers filled with a few centimeters of water (renewed regularly).

### BTM growth rate

Fifteen days later (July 31^st^), we haphazardly collected up to five L3 BTM larvae per box tree (only 6% of plants hosted less than five larvae). We kept them in Petri dishes without food for 24h to make larvae empty their gut and weighed them to the closest 10 µg. In some Petri dishes, we observed cases of cannibalism such that in some instances we could only weight two larvae (Schillé and Kadiri, *personal observation*). For each plant, we therefore calculated the average weight of a L3 larva, dividing the total mass by the number of larvae. Because we did not record the day every single egg hatched, we could not quantify the number of days caterpillars could feed and therefore simply analysed the average weight of a L3 larva.

Larvae were allowed to complete their development on the potted box trees. After every larvae pupated, we counted the number of chrysalis per box tree and weighted them to the closest 10 µg.

### Analyses

All analyses were run in R using libraries nlme and car (Fox et al., 2016; Team, 2018; Pinheiro et al., 2020).

We first looked for spatial patterns in female BTM oviposition. We ran a generalized least square model (GLS) testing the effect of potted tree location in the experimental design (through their *x* and *y* coordinates, **Figure 2)** on the number of clutches per plant (*log*-transformed) from which we explored the associated variogram using the functions *gls* and *Variogram* in the *nlme* library. There was evidence that oviposition was spatially structured, with strong spatial autocorrelation between 1 and 3m **(Figure S1)**.

We tested the effect of prior herbivory on female BTM oviposition (*log*-transformed number of egg clutches) while controlling for spatial non-independence using two independent sets of GLS models. In the first one, we considered prior herbivory as a two-levels factor (attacked vs non-attacked) and used the full data set, whereas in the second one, we treated herbivory as a continuous variable, excluding data from the control treatment. In both cases, we had no particular hypothesis regarding the shape of the spatial correlation structure. We therefore ran separate models with different spatial correlation structures (namely, exponential, Gaussian, spherical, linear and rational quadratic), and compared them based on their AIC (Zuur, 2009). For each model, we computed the Δ*AICc* (*i*.*e*., Δ*i*) as the difference between the AIC of each model *i* and that of the model with the lowest AIC (Burnham & Anderson, 2002). We report and interpret the results of the model with the lowest AIC (see *Results*).

We then tested the effect of prior herbivory on BTM performance using a two-steps approach. We first used two separate ordinary least square models, with the mean weight of L3 larvae (*log*-transformed) or the mean weight of chrysalis (untransformed) as a response variable, the herbivory treatment (non-attacked *vs* attacked) as a two-levels factor and the number of egg clutches as a covariate. Then, we restricted the analyses to plants from the herbivory treatment to test the effect of the percentage of prior herbivory, number of egg clutches and their interaction on the mean weight of L3 larvae (*log*-transformed) and chrysalis, separately. We deleted non-significant interactions prior to the estimation of model coefficient parameters. Finally, we tested the correlation between mean BTM larval weight and mean BTM chrysalis weight at the plant level using Pearson’s correlation.

## Results

We counted a total of 818 egg clutches and 593 larvae on 117 out of 121 plants (*i*.*e*. 96.7%). We counted eggs in 93.4% of plants in the control (non attacked) groups, and in 100% of plants in the herbivory treatment. At individual plant level, the number of egg clutches varied from 0 to 25 (mean ± SD: 6.76 ± 5.11, **Figure 3**).

**Figure 3:**
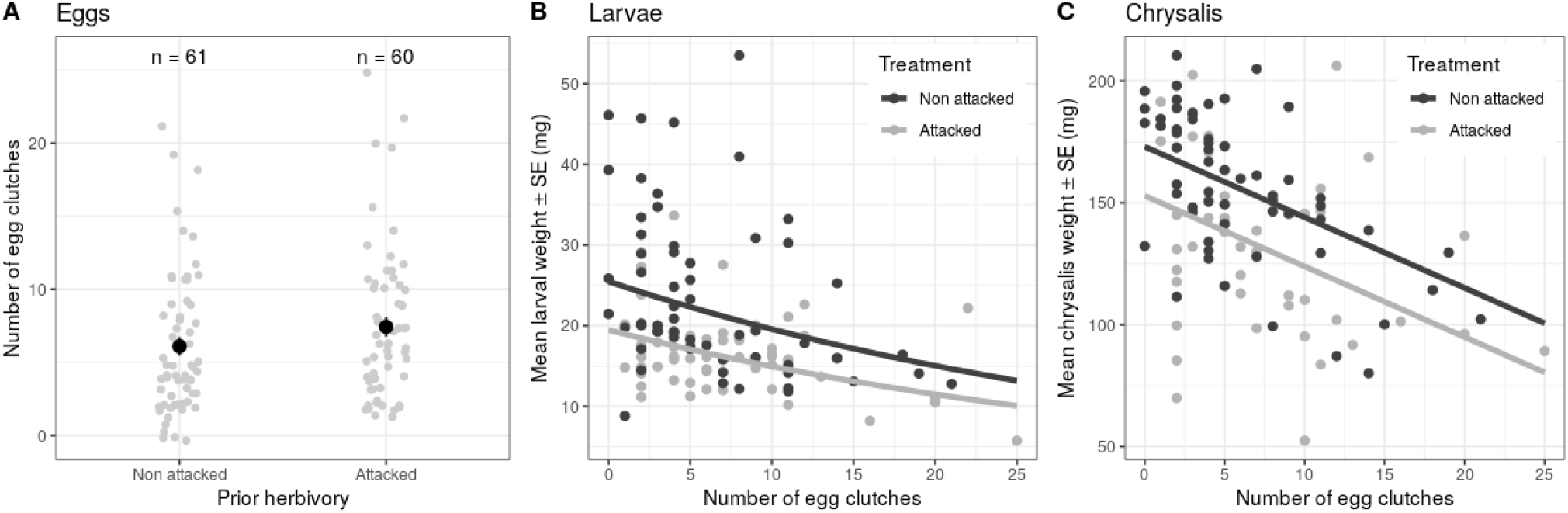
Effects of prior herbivory and conspecific density on (A) the number of egg clutches, (B) L3 larva weight and (C) chrysalis weight. In A, grey dots represent raw data. Black dots and vertical bars represent raw means (± SE). In B and C, dots represent raw data. Black and grey curves represent model predictions for control and herbivory treatments, respectively.

When modelling the effect of prior herbivory on the number of egg clutches using the full data set, the best model (*i*.*e*., model 5 with Δ_i_ = 0, **Table 1**) was the model with a rational quadratic spatial correlation. It was competing with three other models with Δ_i_ < 2 **(Table 1)**. When the analysis excluded data from control plants, the best model was that with a Gaussian spatial correlation **(Table 1.1)**. It was competing with three other models, including that with a rational quadratic spatial correlation (Δ_i_ = 0.2). For the sake of consistency, we therefore used this spatial correlation in further analyses, for it was common to the two analyses. The results were comparable with other spatial correlation structures.

**Table 1:**
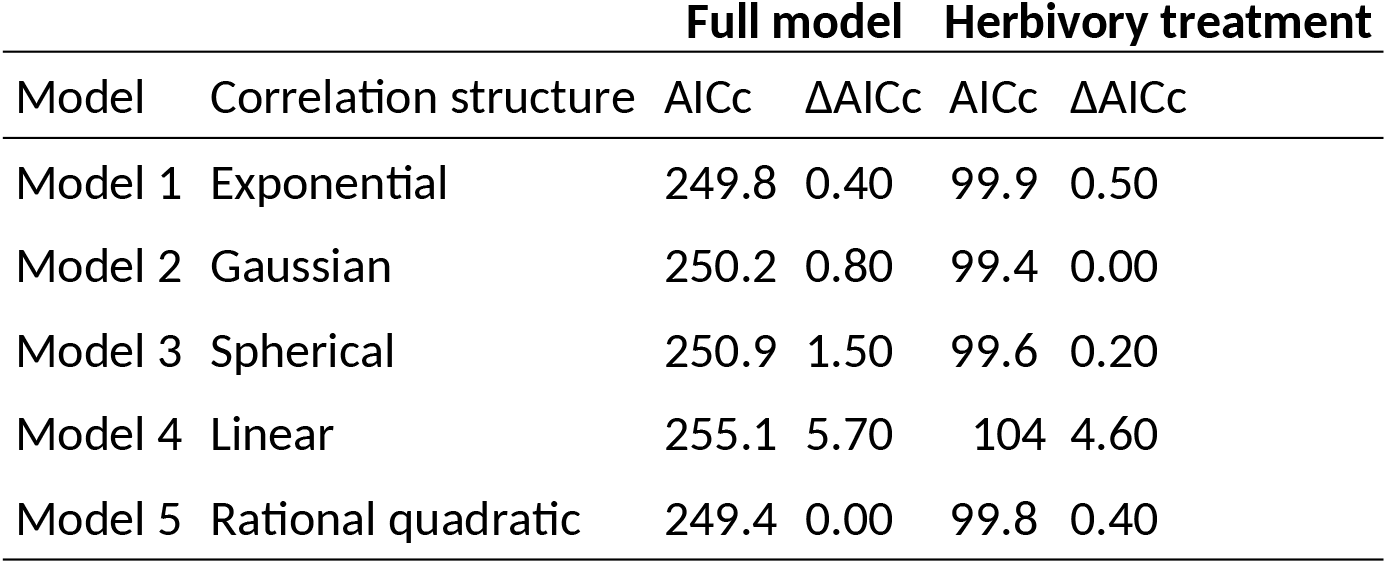
Summary of AIC of GLS models testing the effect of prior herbivory on the number of egg clutches with different spatial correlation structures, for the full dataset and the data set excluding plants from the control treatment.

Herbivory had no significant effect on the number of egg clutches per plant, regardless of whether it was treated as a categorical (model 5, full data set: *F*_1,119_ = 2.91, *P* = 0.09, **Figure 3A**) or continuous variable (model 5, herbivory treatment only: *F*_1, 53_ = 0.8, *P* = 0.374).

The mean weight of BTM larvae varied from 6 to 54 mg (mean ± SD: 20 ± 9 mg). There was a significant, negative relationship between the number of egg clutches on a box tree and subsequent larval weight (**Table 2, Figure 3B**), suggesting intraspecific competition for food. BTM larval weight was lower on box trees that had been previously defoliated (**Table 2, Figure 3B**), regardless of the amount of herbivory (**Table 2**). Larval weight was not significantly affected by the interaction between the herbivory treatment and the number of egg clutches, indicating that intraspecific competition was independent of prior herbivory (**Table 2**). The results were the same regardless of whether herbivory was treated as a categorical or continuous variable (**Table 2**).

**Table 2:**
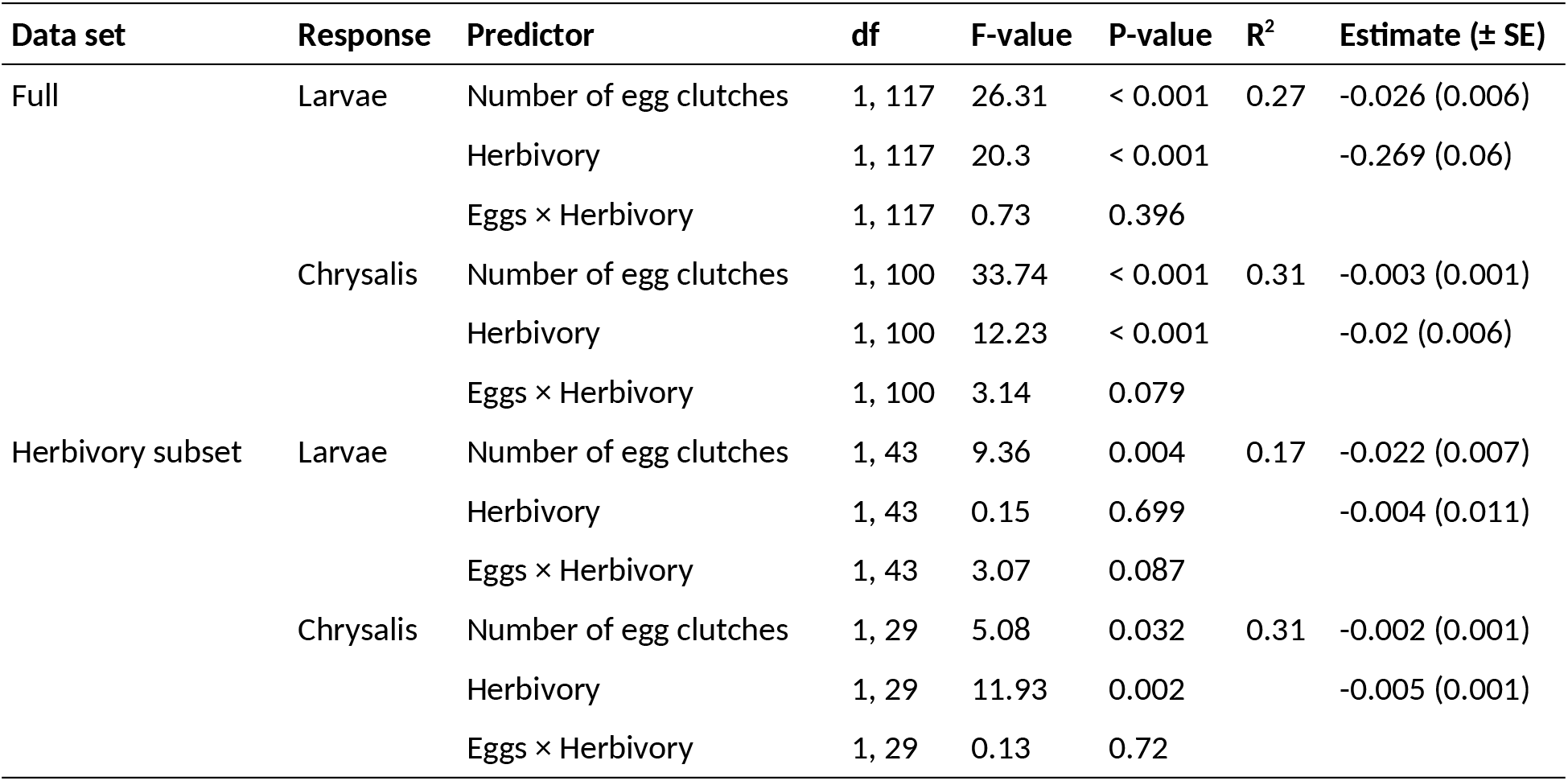
Summary of models testing the effect of prior herbivory (with the full data set or the data set restricted to the herbivory treatment) and initial egg clutch density on mean BTM larvae and chrysalis weight.

### Full model Herbivory treatment

The mean weight of BTM chrysalis varied from 52 to 210 mg (mean ± SD: 145 ± 35 mg, *n* 104). There was a significant positive correlation between the mean weight of BTM larvae and the mean weight of chrysalis (Pearson’s *r* = 0.34, *t*-value = 3.67, *P*-value = < 0.001). The effects of herbivory treatment and number of egg clutches on mean chrysalis weight were very comparable to those observed for BTM larvae: BTM chrysalis weight was lower on box trees that had been previously defoliated (**Table 2, Figure 3C**), and this effect strengthened with an increasing amount of herbivory. There was a significant, negative relationship between the number of egg clutches on a box tree and the subsequent chrysalis weight, which was not significantly affected by the interaction between the herbivory treatment and the number of egg clutches (**Table 2, Figure 3C**).

## Discussion

Our findings reveal that early season herbivory reduces the performance of conspecific individuals that subsequently attack the same host plant later in the plant growing season. This time-lagged intraspecific competition results from a mismatch between female oviposition preference and the performance of its offspring.

### Prior herbivory had no effect BTM oviposition choice

Two possible mechanisms can explain this observation: prior herbivory may have had no effect on box tree characteristics, or female BTM may have been indifferent to them at the time we conducted the experiment.

The first explanation seems unlikely as we found clear evidence that prior herbivory reduced the performance of BTM larvae later in the season. This is fully in line with the numerous studies that have established that insect herbivory induces changes in plant physical and chemical traits, which have profound consequences on herbivores or herbivory on the same host plant later in the season (Wise & Weinberg, 2002; Poelman et al., 2008; Stam et al., 2014; Abdala‐ Roberts et al., 2019; Visakorpi et al., 2019). Therefore, initialRoberts et al., 2019; but see Visakorpi et al., 2019). We cannot dismiss the second explanation that BTM females were indifferent to box tree cues related to earlier herbivory. This may be particularly true in species whose females individually lay several hundred eggs, for which spreading eggs among several host plants may be an optimal strategy (Root & Kareiva, 1984; Hopper, 1999). Consistently, Leuthardt and Baur (2013) observed that BTM females evenly distributed egg clutches among leaves and branches, and that oviposition preference was not dictated by the size of the leaves. Assuming that this behavior is reproducible, the close distance between box-trees that we used in the present experiment (40 cm) could explain the lack of effect of initial defoliation on BTM oviposition behavior. In addition, Leuthard *et al*. (2013) showed that BTM larvae are able to store or metabolise highly toxic alkaloid present in box tree leaves. Even if prior herbivory induced the production of chemical defenses, it is possible that this did not exert any particular pressure upon females for choosing undefended leaves or plants on which to oviposit, as their offspring would have been able to cope with it. Last, BTM larvae proved to be unable to distinguish between box tree leaves infected or not by the box rust *Puccinia buxi*, while their growth is reduced in the presence of the pathogenic fungus (Baur et al., 2019). Altogether, these results suggest that BTM female moths are not influenced by the amount of intact leaves and probably not either by their chemical quality when choosing the host plant, perhaps because of their strong ability to develop on toxic plants. It remains however possible that BTM adults use other cues to select their hosts, such as the presence of conspecific eggs, larvae or chrysalis.

### Prior box tree defoliation by the spring generation of BTM larvae reduced the performance of the next generation

Two alternative, non-mutually exclusive mechanisms can explain this phenomenon. First, the reduced performance of individuals of the second generation can have resulted from induced plant defenses. This explanation is in line with studies that have documented in several plant species reduced herbivore performance and changes in plant-associated herbivore communities linked to induced defenses after prior herbivory (Nykänen & Koricheva, 2004; Karban, 2011; Stam et al., 2014). In the case of multivoltine species, negative relationship between prior herbivory and subsequent larva growth rate could indicate intraspecific plant-mediated cross-generation competition between cohorts of herbivores separated in time (Barnes & Murphy, 2018), which could influence herbivore population dynamics and distribution across host individuals. However, BTM is thought to have broad tolerance to variability in host traits, as suggested by previous observations that BTM larva growth rate did not differ significantly among box-tree varieties (Leuthardt et al., 2013). It is unknown whether herbivory induced changes in host traits are of the same order of magnitude as trait variability among varieties. Assuming variability among varieties is greater, this result goes against the view that reduced performance of larvae of the summer generation resulted from box tree response to prior herbivory. Secondly, reduced performance on previously defoliated plants may partly result from food shortage and increased exploitative competition among larvae of the same cohort. Although free living mandibulate herbivores were described to be less sensitive to competition (Denno et al., 1995), the effect of food shortage may have been exacerbated by the small size of box trees and exploitative competition (Kaygin & Taşdeler, 2019). This explanation is further supported by the fact chrysalis weight was more reduced in plants that were more defoliated by the spring generation of BTM larvae.

### The number of egg clutches laid by BTM female moths correlated negatively with subsequent growth of BTM larvae

This suggests the existence of intraspecific competition for food within the same cohort. Such competition has already been reported, particularly in leaf-miners (Bultman & Faeth, 1986; Faeth, 1992), which are endophagous insect herbivores whose inability to move across leaves makes them particularly sensitive to the choice of oviposition sites by gravid female. In our study, we prevented larvae from moving from one plant to another and noticed that some box trees were completely defoliated by the end of the experiment. Although we did not record this information, it is very likely that larvae first ran out of food in plants on which several egg clutches were laid. We are however unable to determine whether the observed intraspecific competition in this cohort was determined by food shortage, or by herbivore-induced changes in resource quality, or both. In addition, we noticed that the number of chrysalis in 32 control plants (out of 61, *i*.*e*. 52%) was greater than the number of larvae, whereas this only happened in only one previously attacked plant (*i*.*e*. 2%). This suggests that in spite of our precautions some larvae could move from attacked to control plants (Table ??). Together with the fact that patterns of chrysalis weight were very similar to patterns of larval weight, these findings can be seen as another argument in favor of larvae escaping from intraspecific competition on previously attacked plants.

Our findings may have profound implications on our understanding of BTM population dynamics. In many Lepidoptera species, all eggs are present in the ovarioles as the adult molt and larva body mass is proportional to fecundity (i.e., ‘capital breeders,’ (Honěk, 1993; Awmack & Leather, 2002)). As a consequence, host plant quality during larval growth and development is a key determinant of individuals fitness (Awmack & Leather, 2002). Although the relationship between plant quality and herbivore fitness may vary among species (Awmack & Leather, 2002; Moreau et al., 2006; Colasurdo et al., 2009), we speculate that herbivory by the first BTM larva generation reduces the fitness of the second BTM generation, and that this effect may be further strengthened when high population density increases intra-specific cross-generational competition (Tammaru & Haukioja, 1996). These cross-generational effects may thus lead to an important role of density dependence population growth.

## Conclusion

Insect herbivory induces changes in the amount and quality of plant resources, which are responsible for interspecific interactions among herbivores, even in herbivores that are separated in space or time (Poelman et al., 2008; Stam et al., 2014). Our experiment provides evidence that insect herbivory also influences the performance of conspecific herbivores through cross-generational competition, which may ultimately control the overall amount of damage that multivoltine herbivore species can cause to plants. Cross-generational competition may increase development time of individuals of the next generation, thereby increasing their vulnerability to natural enemies [the *slow-growth-high-mortality hypothesis*; Coley et al. (2006); Benrey & Denno (1997); Uesugi (2015)]. If this is the case, on the one hand stronger top-down control can be exerted on herbivores feeding on previously attacked hosts, which could reduce the overall amount of damage to the host plant. On the other hand, if herbivores take longer to develop, they may cause more damage to plants, in particular to those with poor nutritional quality, due to compensatory feeding (Simpson & Simpson, 1990; Milanovic et al., 2014). Our results highlight the overlooked ecological importance of time-lagged intraspecific competition (Barnes & Murphy, 2018). In the face of global warming, which shortens the generation time of many insect herbivores and thus increases voltinism (Jactel et al., 2019), it is particularly necessary to elucidate the consequences of cross-generational interactions on the population dynamics of multivoltine herbivore species.

## Data accessibility

Raw data as well as codes of statistic analysis are available in supplementary material and on the Inrae dataverse: Castagneyrol, Bastien; van Halder, Inge; Kadiri, Yasmine; Schillé, Laura; Jactel, Hervé, 2020, “Raw data for the paper ‘Host-mediated, cross-generational intraspecific competition in a herbivore species,’ https://doi.org/10.15454/KMUX39, Portail Data INRAE, V3.0.

## Supplementary material

Script and codes are available online Inrae dataverse: Castagneyrol, Bastien; van Halder, Inge; Kadiri, Yasmine; Schillé, Laura; Jactel, Hervé, 2020, “Raw data for the paper ‘Host-mediated, cross-generational intraspecific competition in a herbivore species,’ https://doi.org/10.15454/KMUX39, Portail Data INRAE, V3.0.

## Acknowledgements

Version 5 of this preprint has been peer-reviewed and recommended by Peer Community In Ecology (https://doi.org/10.24072/pci.ecology.100072). We thank Alex Stemmelen, and Yannick Mellerin for their help in BTM rearing and data collection. This research was founded by the HOMED project, which received funding from the European Union’s Horizon 2020 research and innovation program under grant agreement No. 771271. We thank Inês Fragata, Raul Costa-Pereira and Sara Magalhães for their helpful comments on earlier version of this manuscript.

## Appendix

Supplementary material, available online: https://doi.org/10.15454/KMUX39

Scripts and codes for statistical analyses: https://doi.org/10.15454/KMUX39

