## Supplementary material for "Host-mediated, cross-generational intraspecific competition in a herbivore species": Castagneyrol_Cydalima_RCodes_Rev2_Manuscript+SM+Reviews.pdf (from another version of this preprint: version 4, 2021-02-11)

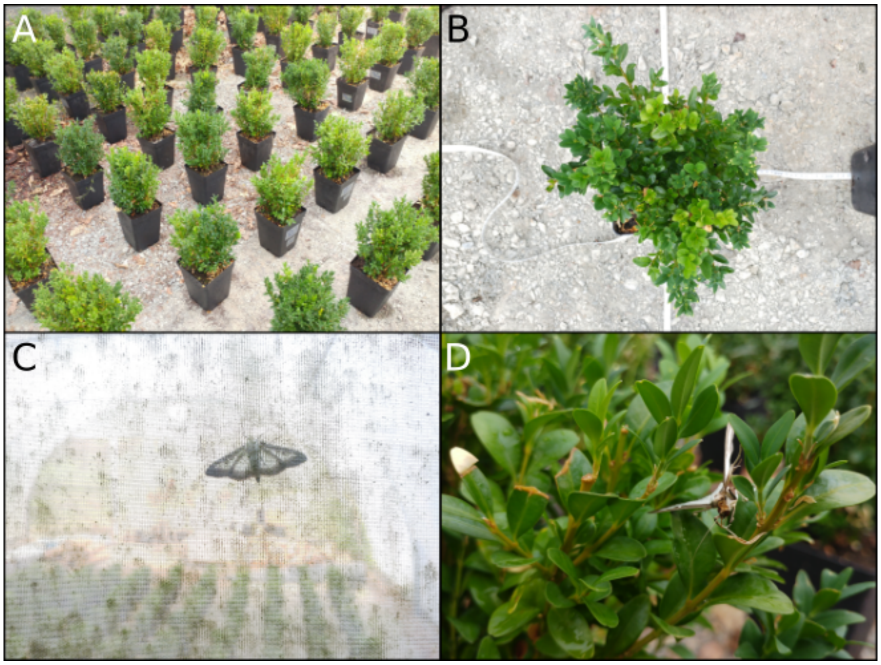


Figure 1: The study design and model species. The two top photos (A, B) illustrate the experimental design and in particular distance among potted plants. Photo C is a view of the greenhouse from the outside, with an adult box tree moth in the foreground, and potted plants in the background. Photo D shows an adult box tree moth on a box tree branch, shortly after it was released.

performance should have been independent of cues emitted by BTM larvae themselves or by their frass (Sato et al., 1999; Molnár et al., 2017) and therefore were only plant-mediated.

###### 1.2.4 BTM host choice

In order to test whether initial defoliation of focal plants influenced host choice for oviposition by BTM females, we counted egg clutches on every branch of every box tree on July $17^{th}$. Once eggs were counted, we moved box trees to another greenhouse. To prevent larvae from moving from one potted plant to another, we installed box trees in plastic saucers filled with a few centimeters of water (renewed regularly).

###### 1.2.5 BTM growth rate

Fifteen days later (July $31^{st}$), we haphazardly collected up to five L3 BTM larvae per box tree (only 6% of plants hosted less than five larvae). We kept them in Petri dishes without food for 24h to make larvae empty their gut and weighed them to the closest 10 µg. In some Petri dishes, we observed cases of cannibalism such that in some instances we could only weight two larvae (Schillé and Kadiri, *personal observation*). For each plant, we therefore calculated the average weight of a L3 larva, dividing the total mass by the number of larvae. Because we did not record the day every single egg hatched, we could not quantify the number of days caterpillars could feed and therefore simply analysed the average weight of a L3 larva.

Larvae were allowed to complete their development on the potted box trees. After every larvae pupated, we counted the number of chrysalis per box tree and weighted them to the closest 10 µg.

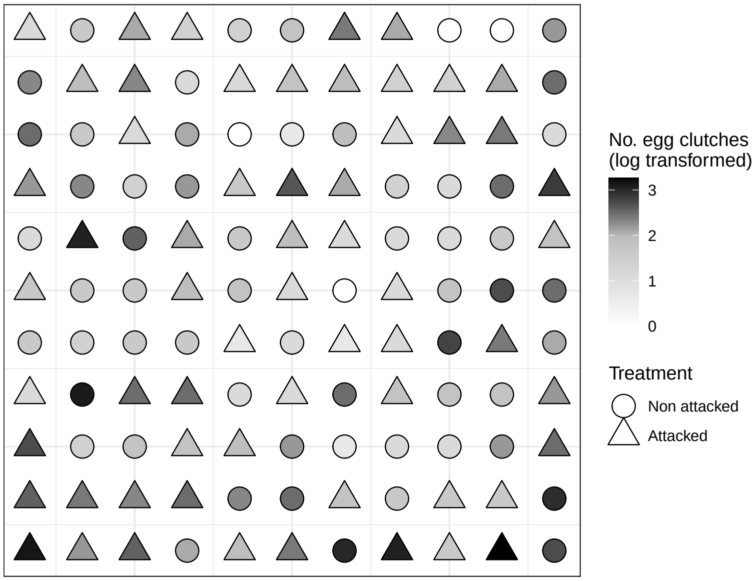


Figure 2: Experimental design. Pots were 40 cm apart. Circles and triangles represent non-attacked (control) and attacked trees. Scale colour represents the number of egg clutches per box tree (log-transformed).

When modelling the effect of prior herbivory on the number of egg clutches using the full data set, the best model (*i.e.*, model 5 with $\Delta_i = 0$, **Table 1**) was the model with a rational quadratic spatial correlation. It was competing with three other models with $\Delta_i < 2$ **(Table 1)**. When the analysis excluded data from control plants, the best model was that with a Gaussian spatial correlation **(Table 1)**. It was competing with three other models, including that with a rational quadratic spatial correlation ($\Delta AIC = 0.2$). For the sake of consistency, we therefore used this spatial correlation in further analyses, for it was common to the two analyses. The results were comparable with other spatial correlation structures.

Table 1: Summary of AIC of GLS models testing the effect of prior herbivory on the number of egg clutches with different spatial correlation structures, for the full dataset and the data set excluding plants from the control treatment.

| | | Full model | | Herbivory treatment | |
|---|---|---|---|---|---|
| Model | Correlation structure | AIC | $\Delta$ | AIC | $\Delta$ |
| Model 1 | Exponential | 249.8 | 0.4 | 99.9 | 0.5 |
| Model 2 | Gaussian | 250.2 | 0.8 | 99.4 | 0.0 |
| Model 3 | Spherical | 250.9 | 1.5 | 99.6 | 0.2 |
| Model 4 | Linear | 255.1 | 5.7 | 104.0 | 4.6 |
| Model 5 | Rational quadratic | 249.4 | 0.0 | 99.8 | 0.4 |

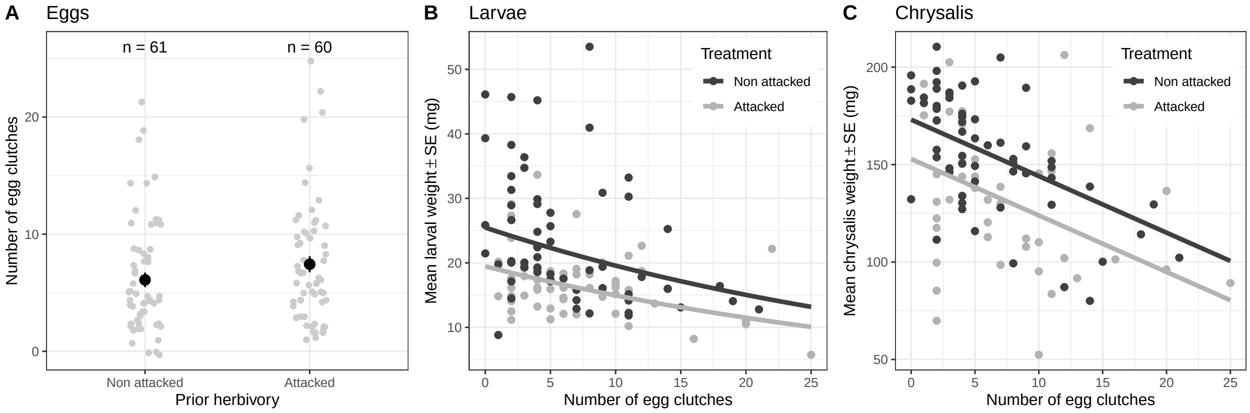


Figure 3: Effects of prior herbivory and conspecific density on (A) the number of egg clutches, (B) L3 larva weight and (C) chrysalis weight. In A, grey dots represent raw data. Black dots and vertical bars represent raw means (+/- SE). In B and C, dots represent raw data. Black and grey curves represent model predictions for control and herbivory treatments, respectively.

Table 2: Summary of models testing the effect of prior herbivory (with the full data set or the data set restricted to the herbivory treatment) and initial egg clutch density on mean BTM larvae and chrysalis weight

| Data set | Response | Predictor | df | F-value | P-value | $R^2$ | Estimate (SE) |
|---|---|---|---|---|---|---|---|
| Full | Larvae | Number of egg clutches | 1, 117 | 26.31 | < 0.001 | 0.27 | -0.026 (0.006) |
| | | Herbivory | 1, 117 | 20.30 | < 0.001 | | -0.269 (0.06) |
| | | Eggs x Herbivory | 1, 117 | 0.73 | 0.396 | | |
| | Chrysalis | Number of egg clutches | 1, 100 | 33.74 | < 0.001 | 0.31 | -0.003 (0.001) |
| | | Herbivory | 1, 100 | 12.23 | < 0.001 | | -0.02 (0.006) |
| | | Eggs x Herbivory | 1, 100 | 3.14 | 0.079 | | |
| Herbivory subset | Larvae | Number of egg clutches | 1, 43 | 9.36 | 0.004 | 0.17 | -0.022 (0.007) |
| | | Herbivory | 1, 43 | 0.15 | 0.699 | | -0.004 (0.011) |
| | | Eggs x Herbivory | 1, 43 | 3.07 | 0.087 | | |
| | Chrysalis | Number of egg clutches | 1, 29 | 5.08 | 0.032 | 0.31 | -0.002 (0.001) |
| | | Herbivory | 1, 29 | 11.93 | 0.002 | | -0.005 (0.001) |
| | | Eggs x Herbivory | 1, 29 | 0.13 | 0.72 | | |

#### 1.8 Conflict of interest

The authors of this preprint declare that they have no financial conflict of interest with the content of this article. Bastien Castagneyrol is one of the *PCI Ecology* recommenders.

### 2 Appendix

#### 2.1 Supplementary material

**Figure S1** - Semivariogram of the number of egg clutches as a function of distance among box trees.

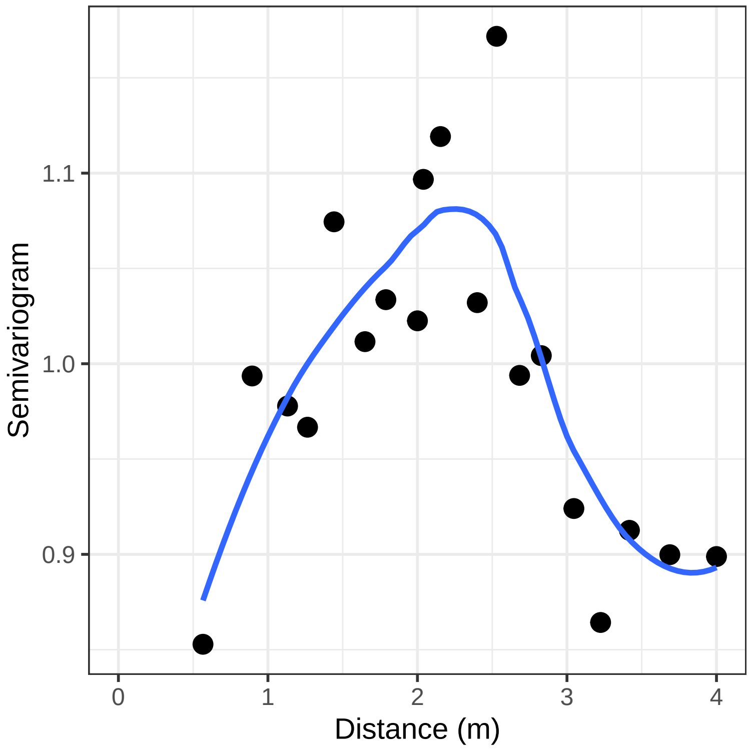

Table 3: Repartition of egg clutches, larvae and chrysalis across box trees with or without prior herbivory. Numbers correspond to mean (± sd) and total number of egg clutches, larvae or chrysalis (n).

| Response variable | Control | Herbivory treatment |
|---|---|---|
| Egg clutches | 6.1 (4.87), n = 372 | 7.43 (5.3), n = 446 |
| Larvae | 4.84 (0.61), n = 295 | 4.97 (0.18), n = 298 |
| Chrysalis | 6.8 (5.78), n = 415 | 1.85 (1.79), n = 111 |

#### 2.2 Raw data

**Table S2** - Raw data used in the present manuscript: `x` and `y` are the position of each box tree in the green house; `Treatment` is the prior herbivory treatment; `Clutch.number` is the total number of egg clutches counted on a given box tree; `N.L3` is the number of retrieved L3 larvae, `L3.mean` is the mean weight of a L3 larvae (g); `N.chrysalids` is the number of retrieved chrysalis; `Chrysalid.mean` is the mean weight of a chrysalis. `Herbivory` is the % of leaves consumed by box tree moth larvae, which was either measured or estimated where raw data was missing (`Herbivory_source`).

| x | y | Treatment | Clutch.number | N.L3 | L3.mean | N.chrysalids | Chrysalid.mean | Herbivory | Herbivory_s |
|---|---|---|---|---|---|---|---|---|---|
| 1 | 1 | Attacked | 22 | 5 | 0.0221740 | 0 | NaN | 7.3 | Estimated |
| 2 | 1 | Attacked | 8 | 5 | 0.0183980 | 0 | NaN | 8.8 | Estimated |
| 3 | 1 | Attacked | 12 | 5 | 0.0187360 | 1 | 0.1020100 | 8.8 | Imputed |
| 5 | 1 | Attacked | 6 | 5 | 0.0146140 | 0 | NaN | 16.4 | Estimated |
| 6 | 1 | Attacked | 10 | 5 | 0.0165620 | 2 | 0.1101750 | 10.5 | Estimated |
| 8 | 1 | Attacked | 20 | 5 | 0.0110140 | 1 | 0.0961900 | 8.7 | Estimated |
| 9 | 1 | Attacked | 4 | 5 | 0.0132300 | 1 | 0.1438500 | 7.4 | Estimated |
| 10 | 1 | Attacked | 25 | 5 | 0.0057520 | 1 | 0.0891900 | 11.7 | Estimated |
| 1 | 2 | Attacked | 12 | 5 | 0.0226500 | 2 | 0.2062350 | 6.2 | Estimated |
| 2 | 2 | Attacked | 10 | 5 | 0.0162200 | 1 | 0.0524100 | 0.0 | Imputed |
| 3 | 2 | Attacked | 9 | 5 | 0.0200760 | 0 | NaN | 4.2 | Estimated |
| 4 | 2 | Attacked | 11 | 5 | 0.0211200 | 5 | 0.1465000 | 4.8 | Estimated |
| 7 | 2 | Attacked | 5 | 5 | 0.0112560 | 0 | NaN | 9.0 | Estimated |
| 9 | 2 | Attacked | 4 | 5 | 0.0161760 | 1 | 0.1338800 | 15.1 | Estimated |
| 10 | 2 | Attacked | 4 | 5 | 0.0173680 | 1 | 0.1706800 | 9.4 | Estimated |
| 1 | 3 | Attacked | 14 | 5 | 0.0159000 | 4 | 0.1686525 | 8.6 | Imputed |
| 4 | 3 | Attacked | 5 | 5 | 0.0159420 | 2 | 0.1380100 | 0.0 | Imputed |
| 5 | 3 | Attacked | 6 | 5 | 0.0121100 | 1 | 0.1319100 | 8.6 | Imputed |
| 11 | 3 | Attacked | 11 | 5 | 0.0101960 | 1 | 0.0836300 | 7.2 | Estimated |
| 1 | 4 | Attacked | 2 | 5 | 0.0111600 | 2 | 0.1224050 | 11.4 | Estimated |
| 3 | 4 | Attacked | 11 | 5 | 0.0157420 | 0 | NaN | 8.6 | Imputed |
| 4 | 4 | Attacked | 11 | 5 | 0.0158140 | 4 | 0.1557575 | 6.8 | Estimated |
| 6 | 4 | Attacked | 2 | 5 | 0.0238660 | 2 | 0.1728600 | 11.7 | Estimated |
| 8 | 4 | Attacked | 5 | 5 | 0.0187260 | 2 | 0.1527050 | 10.0 | Estimated |
| 11 | 4 | Attacked | 8 | 5 | 0.0181900 | 0 | NaN | 10.3 | Estimated |
| 5 | 5 | Attacked | 1 | 5 | 0.0201820 | 1 | 0.1914500 | 2.7 | Estimated |
| 7 | 5 | Attacked | 1 | 5 | 0.0148320 | 3 | 0.1752800 | 8.9 | Estimated |
| 8 | 5 | Attacked | 2 | 5 | 0.0176160 | 1 | 0.0853400 | 9.1 | Estimated |
| 10 | 5 | Attacked | 10 | 4 | 0.0171925 | 1 | 0.1453700 | 9.0 | Estimated |
| 1 | 6 | Attacked | 4 | 5 | 0.0158160 | 2 | 0.1774000 | 2.2 | Estimated |
| 4 | 6 | Attacked | 6 | 5 | 0.0161500 | 0 | NaN | 6.8 | Estimated |
| 6 | 6 | Attacked | 2 | 4 | 0.0148600 | 0 | NaN | 10.9 | Estimated |
| 8 | 6 | Attacked | 2 | 5 | 0.0273120 | 2 | 0.0698850 | 17.0 | Estimated |
| 2 | 7 | Attacked | 20 | 5 | 0.0104900 | 4 | 0.1364600 | 3.6 | Estimated |
| 4 | 7 | Attacked | 7 | 5 | 0.0275520 | 5 | 0.1299800 | 12.5 | Estimated |
| 6 | 7 | Attacked | 6 | 5 | 0.0143660 | 2 | 0.1127850 | 9.7 | Estimated |
| 7 | 7 | Attacked | 2 | 5 | 0.0145880 | 0 | NaN | 17.2 | Estimated |
| 11 | 7 | Attacked | 5 | 5 | 0.0129260 | 3 | 0.1438500 | 9.5 | Estimated |
| 1 | 8 | Attacked | 8 | 5 | 0.0161140 | 0 | NaN | 9.5 | Estimated |
| 5 | 8 | Attacked | 4 | 5 | 0.0336620 | 4 | 0.1512050 | 0.0 | Imputed |
| 6 | 8 | Attacked | 13 | 5 | 0.0136940 | 1 | 0.0916800 | 5.6 | Imputed |
| 7 | 8 | Attacked | 7 | 5 | 0.0119960 | 0 | NaN | 7.6 | Estimated |
| 11 | 8 | Attacked | 16 | 5 | 0.0082180 | 5 | 0.1013240 | 5.6 | Imputed |
| 3 | 9 | Attacked | 2 | 5 | 0.0124840 | 2 | 0.1309350 | 0.0 | Imputed |
| 8 | 9 | Attacked | 2 | 5 | 0.0140740 | 1 | 0.0996800 | 5.6 | Imputed |
| 9 | 9 | Attacked | 9 | 5 | 0.0147260 | 3 | 0.1120367 | 15.1 | Estimated |
| 10 | 9 | Attacked | 10 | 5 | 0.0121140 | 3 | 0.1454233 | 7.9 | Estimated |
| 2 | 10 | Attacked | 6 | 5 | 0.0183400 | 3 | 0.1203367 | 0.0 | Imputed |
| 3 | 10 | Attacked | 9 | 5 | 0.0159820 | 3 | 0.1078233 | 6.8 | Estimated |
| 5 | 10 | Attacked | 2 | 5 | 0.0291080 | 2 | 0.1450000 | 10.5 | Estimated |
| 6 | 10 | Attacked | 5 | 5 | 0.0185740 | 0 | NaN | 8.6 | Estimated |
| 7 | 10 | Attacked | 6 | 5 | 0.0177680 | 0 | NaN | 12.4 | Estimated |
| 8 | 10 | Attacked | 3 | 5 | 0.0149260 | 1 | 0.2025200 | 9.4 | Estimated |
| 9 | 10 | Attacked | 3 | 5 | 0.0195980 | 2 | 0.1319950 | 8.2 | Estimated |
| 10 | 10 | Attacked | 7 | 5 | 0.0157780 | 2 | 0.0985400 | 16.2 | Estimated |
| 1 | 11 | Attacked | 2 | 5 | 0.0161540 | 5 | 0.1175720 | 8.6 | Imputed |
| 3 | 11 | Attacked | 7 | 5 | 0.0190760 | 5 | 0.1385900 | 8.5 | Estimated |
| [illegible] | [illegible] | [illegible] | [illegible] | [illegible] | [illegible] | [illegible] | [illegible] | [illegible] | [illegible] |

#### 2.3 R codes used to generate this report

```r
library(tidyverse)
library(knitr)
library(kableExtra)
library(nlme)
library(car)
library(here)

wd = here()

my.ggplot <- function(){
  theme_bw() +  theme(legend.key = element_blank())
}

knitr::opts_chunk$set(echo= F, warning= F, message = F, results = "hide",
                      fig.width=4, fig.height=4, dpi= 100, fig.pos = "H",
                      fig.path = paste0(wd, '/Figures/'),
                      output.dir = paste0(wd, '/Outputs/'))

d = read.csv(paste0(wd, '/Data/Castagneyrol_cydalima_data.csv'), header = T)

str(d)

d =
  d %>% mutate(L3.mean = Weight.L3 / N.L3) %>%
  mutate(Treatment = factor(Treatment, levels = c("Non attacked", "Attacked")))

knitr::include_graphics(paste0(wd, '/Figures/Figure 1.png'))

d %>%
  ggplot(aes(x, y, shape = Treatment, fill = log1p(Clutch.number))) +
  # geom_rect(aes(xmin = 1.5, ymin = 1.5, xmax = 10.5, ymax = 10.5), fill = 'grey90') +
  geom_point(size = 6) +
  scale_shape_manual(values = c(21, 24)) +
  my.ggplot() +
  labs(x = "", y = "") +
  scale_fill_gradientn(colours = c('white', 'grey', 'black'),
                       values = c(0,0.6,1), name = 'No. egg clutches\n(log transformed)') +
  theme(axis.ticks.x = element_blank(),
        axis.text.x = element_blank(),
        axis.ticks.y = element_blank(),
        axis.text.y = element_blank())
d.center = droplevels(d[d$x > 1 & d$x < 11 & d$y > 1 & d$y < 11,])
m1a = gls(log1p(Clutch.number) ~ Treatment,
          correlation = corExp(form = ~x + y, nugget = TRUE),
          data = d, na.action = "na.omit")
m1b = gls(log1p(Clutch.number) ~ Treatment,
          correlation = corGaus(form = ~x + y, nugget = TRUE),
          data = d, na.action = "na.omit")
m1c = gls(log1p(Clutch.number) ~ Treatment,
          correlation = corSpher(form = ~x + y, nugget = TRUE),
          data = d, na.action = "na.omit")
```

```r
m1d = gls(log1p(Clutch.number) ~ Treatment,
          correlation = corLin(form = ~x + y, nugget = TRUE),
          data = d, na.action = "na.omit")
m1e = gls(log1p(Clutch.number) ~ Treatment,
          correlation = corRatio(form = ~x + y, nugget = TRUE),
          data = d, na.action = "na.omit")

l1 = list(m1a, m1b, m1c, m1d, m1e)

l2 = list(
m1a_2 = update(m1a, ~ Herbivory,
               data = d[d$Herbivory > 0 &
                          d$Herbivory_source == "Estimated",]),
m1b_2 = update(m1b, ~ Herbivory,
               data = d[d$Herbivory > 0 &
                          d$Herbivory_source == "Estimated",]),
m1c_2 = update(m1c, ~ Herbivory,
               data = d[d$Herbivory > 0 &
                          d$Herbivory_source == "Estimated",]),
m1d_2 = update(m1d, ~ Herbivory,
               data = d[d$Herbivory > 0 &
                          d$Herbivory_source == "Estimated",]),
m1e_2 = update(m1e, ~ Herbivory,
               data = d[d$Herbivory > 0 &
                          d$Herbivory_source == "Estimated",])
)

AIC_1 = round(unlist(lapply(l1, function(x){AIC(x)})),1)
delta_1 = AIC_1 - min(AIC_1)
AIC_2 = round(unlist(lapply(l2, function(x){AIC(x)})),1)
delta_2 = AIC_2 - min(AIC_2)

Table_AIC = data.frame(AIC_1, delta_1, AIC_2, delta_2)

plot.resid = function(m){
  df = data.frame(f = fitted(m), r = residuals(m))
  A = df %>%
    ggplot(aes(f, r)) + my.ggplot() + geom_point() +
    geom_smooth(method = "lm", se = F) + geom_hline(yintercept = 0)
  B = df %>%
    ggplot(aes(r)) + my.ggplot() + geom_histogram()
  cowplot::plot_grid(A,B)
}


m_larvae = lm(log(L3.mean) ~ Clutch.number * Treatment , d)
m_larvae2 = lm(log(L3.mean) ~ Clutch.number * Herbivory,
               d[d$Herbivory > 0 &
                   d$Herbivory_source == "Estimated",])
# plot.resid(m_larvae2)

d =
  d %>%
```

```
  mutate(Chrysalid.mean = Weight.chrysalids / N.weighted.chysalids)

m_chrys = lm(Chrysalid.mean ~ Clutch.number * Treatment , d)
m_chrys2 = lm(Chrysalid.mean ~ Clutch.number * Herbivory,
              d[d$Herbivory > 0  &
                  d$Herbivory_source == "Estimated",])

#plot(m_chrys2)

Anova(glm(Clutch.01 ~ Treatment, d %>%  mutate(Clutch.01 = ifelse(Clutch.number == 0, 0, 1)), family =


# lapply(l1, function(x) anova(x))
# lapply(l2, function(x) anova(x))
Table_AIC %>%
  remove_rownames() %>%
  mutate(Model = paste('Model', 1:5),
         'Correlation structure' = c('Exponential', 'Gaussian', 'Spherical',
                                     'Linear', 'Rational quadratic')) %>%
  select(Model, `Correlation structure`, everything()) %>%
  kable(col.names = c("Model", "Correlation structure", "AIC", "$\\Delta$", "AIC", "$\\Delta$"),
        caption = "Summary of AIC of GLS models testing the effect of prior
        herbivory on the number of egg clutches with different spatial correlation
        structures, for the full dataset and the data set excluding plants from the
        control treatment.") %>%
  kableExtra::kable_styling() %>%
  add_header_above(c(" " = 2, "Full model" = 2, "Herbivory treatment" = 2))
anova(m1e)

Fig_3A =
  d %>%
  ggplot(aes(Treatment, Clutch.number)) +
  my.ggplot() +
  geom_point(position = position_jitter(0.1), colour = "grey80") +
  stat_summary(size = 0.6) +
  labs(x = "Prior herbivory", y = "Number of egg clutches") +
  annotate(geom = 'text', x = 1:2, y = 26, label = paste("n =", c(61, 60)))
res_larvae = anova(m_larvae, test = "m")
n = expand.grid(Treatment = levels(d$Treatment), Clutch.number = seq(0, 25))
p = predict(update(m_larvae, ~.- Clutch.number:Treatment), newdata = n)
n$Fit = 1000 * exp(p)

Fig_3B =
  d %>%
  ggplot(aes(Clutch.number, 1000 * L3.mean, colour = Treatment)) +
  my.ggplot() +
  geom_point(size = 2) +
  scale_colour_manual(values = c("grey25", "grey70")) +
  labs(x = "Number of egg clutches",
       y = expression("Mean larval weight" %+-% "SE (mg)")) +
  geom_line(data = n, aes(Clutch.number, Fit), size = 1.5) +
  theme(legend.position = c(0.8, 0.85))
r = with(d, cor.test(Chrysalid.mean, L3.mean))
```

```r
r_corr = round(r$estimate,2)
r_tval = round(r$statistic,2)
r_pval = ifelse(round(r$p.value,3) < 0.001, "< 0.001", round(r$p.value,3))
res_chrys = anova(m_chrys)
n = expand.grid(Treatment = levels(d$Treatment), Clutch.number = seq(0, 25))
p = predict(update(m_chrys, ~.- Clutch.number:Treatment), newdata = n)
n$Fit = 1000 * p

Fig_3C =
  d %>%
  ggplot(aes(Clutch.number, 1000 * Chrysalid.mean, colour = Treatment)) +
  my.ggplot() +
  geom_point(size = 2) +
  scale_colour_manual(values = c("grey25", "grey70")) +
  labs(x = "Number of egg clutches",
       y = expression("Mean chrysalis weight" %+-% "SE (mg)")) +
  geom_line(data = n, aes(Clutch.number, Fit), size = 1.5) +
  theme(legend.position = c(0.8, 0.85))

cowplot::plot_grid(
  Fig_3A + labs(title = 'Eggs'),
  Fig_3B + labs(title = 'Larvae'),
  Fig_3C + labs(title = 'Chrysalis'),
  ncol = 3,
  labels = c('A', 'B', 'C')
)
f = function(model, response, data_set){
  ANOVA = anova(model, test = "m")
  Fval = function(ANOVA) {round(ANOVA[,4], 2)}
  Pval = function(ANOVA) {ifelse(ANOVA[,5] < 0.001, '< 0.001', round(ANOVA[,5], 3))}

  if(Pval(ANOVA)[3] < 0.05){
    b = round(summary(model)$coefficients[-1,1], 3)
    b_se = round(summary(model)$coefficients[-1,2], 3)
    Estimate = paste(b, ' (', b_se, ')', sep = '')

    }else{

    form = formula(paste("~",
                         paste(attr(model$terms, "variables")[[3]],
                               attr(model$terms, "variables")[[4]],
                               sep ="+")))
    b = round(summary(update(model, formula. = form))$coefficients[-1,1], 3)
    b_se = round(summary(update(model, formula. = form))$coefficients[-1,2], 3)
    Estimate = c(paste(b, ' (', b_se, ')', sep = ''), '')
  }

  return(df =
           data.frame(
             Data = c(data_set, "", ""),
             Response = c(response, '', ''),
             Predictor = c("Number of egg clutches", "Herbivory", "Eggs x Herbivory"),
             df = paste(ANOVA$Df[1:3], rep(ANOVA$Df[4], 3), sep = ", "),
```

```
             `F-value` = Fval(ANOVA)[-4],
             `P-value` = Pval(ANOVA)[-4],
             R2 = c(round(summary(model)$adj.r.squared, 2), "", ""),
             Estimate = Estimate))
}


rbind(
  f(m_larvae, response = "Larvae", data_set = "Full"),
  f(m_chrys, response = "Chrysalis", data_set = ""),
  f(m_larvae2, response = "Larvae", data_set = "Herbivory subset"),
  f(m_chrys2, response = "Chrysalis", data_set = "")) %>%
  kable(caption = "Summary of models testing the effect of prior herbivory
        (with the full data set or the data set restricted to the herbivory
        treatment) and initial egg clutch density on mean BTM larvae and
        chrysalis weight",
        col.names = c("Data set", "Response", "Predictor", "df", "F-value",
                      "P-value", "R²", "Estimate (SE)"),
        escape = T, digit = 2) %>%
  kable_styling() %>%
  collapse_rows(columns = 1:2, valign = "top")


m0 = gls(log1p(Clutch.number) ~ x + y, data = d, na.action = "na.omit")
variog0 <- Variogram(m0, form = ~x + y, resType = "pearson", nugget = T)
variog0 %>%
  ggplot(aes(0.4*dist, variog)) +
  my.ggplot() +
  geom_point(size = 3) +
  geom_smooth(se = F) +
  labs(x = "Distance (m)", y = "Semivariogram") +
  xlim(0, 4)
d %>%
  select(Treatment, Clutch.number, N.L3, N.chysalids) %>%
  gather(Resp, Val, 2:4) %>%
  group_by(Treatment, Resp) %>%
  summarise(N = sum(Val), M = round(mean(Val),2), SD = round(sd(Val),2)) %>%
  mutate(Val = paste(M, ' (', SD, ')', ', n = ', N, sep = '')) %>%
  select(- N, - M, - SD) %>%
  mutate(Resp = factor(Resp, levels = c('Clutch.number', 'N.L3', 'N.chysalids'),
                       labels = c('Egg clutches', 'Larvae', 'Chrysalis'))) %>%
  spread(Treatment, Val) %>%
  kable(col.names = c('Response variable', 'Control', 'Herbivory treatment'),
        caption = "Repartition of egg clutches, larvae and chrysalis across box trees with or without pr
  kable_styling()
d %>%
  rename(N.chrysalids = N.chysalids) %>%
  select(x, y, Treatment, Clutch.number, N.L3, L3.mean, N.chrysalids, Chrysalid.mean, Herbivory, Herbiv
  kable() %>% kable_styling()
```

### 3 Responses to reviewers' comments

#### 3.1 First round

**Dear Dr Magalhães,**

**We would like to thank you for your constructive and helpful comments. We revised the original manuscript accordingly. Significant changes in the manuscript are highlighted in bold characters. You may find our responses to your comments and to the two reviewers below, highlighted with bold characters[1]. Wherever only minor changes were requested, we simply indicated "[R] — Done", otherwise, we justified what we changed, or not, in the manuscript.**

**We hope that the revised version of our manuscript has addressed every concerns and will be suitable for recommendation**.

**Best regards,**

**Bastien Castagneyrol, on behalf of co-authors**.

---

Dear authors,

First of all, I deeply apologize for having taken so long to comment on this manuscript. I hope that the quality of the reviews compensates for this long wait... I found this article interesting and straightforward. I particularly appreciated the scale and nature of the experiment, being an intermediate between a lab and a field experiment. The thorough and insightful comments of the two reviewers also point in the same direction: they both enjoyed the manuscript very much. They do, however, suggest a number of changes that I think should be addressed in the revised version of the manuscript. In particular, they both suggest including more variables in your analysis, which they (and I) believe you can do based on the data you already have. If that is the case, I think it is a cost-effective means to make your article more complete.

**[Response] — We followed recommendations made by the two reviewers and yourself and provide below detailed answers wherever necessary**

Below I place my own comments, and I would be happy to look at a revised version of this paper soon. I only have two relatively major and a lot of minor comments.

1. I agree with one of the reviewers in that the introduction could be a bit further streamlined. If I understand correctly, the first paragraph is about preference-performance correlations, the second about preference, and the third a bit about performance and then another bit about preference-performance. You also go a bit back and forth concerning the effects of conspecifics and that of heterospecifics. I don't want to impose my view on the introduction of a paper that is not mine, but I would like to feel that, whatever the structure chosen, it is apparent to the reader. In any case, I would exchange the order of second and third paragraph, first differences in performance and then oviposition choice, because the latter does not make sense without the former and the reverse is not true.

**[R] — We changed the order of the two paragraphs as suggested and modified several sentences to improve the flow. Please note that we did not highlight every single change in the manuscript, only the most important additions.**

2. I think the reader needs some information on how larval weight correlates (or may correlate) with fitness in this (or related) species. This would allow discriminating among the two main possible interpretations for this data set, namely (a) they don't discriminate because the consequences for fitness are not strong enough or (b) they don't discriminate because they don't have access to reliable cues.

**[R] — We now discuss this question:**

[1] we have removed the bold characters after the first round of review round to make the article easier to read.

*Our findings may have profound consequences on BTM population dynamics. In most of Lepidoptera species, all the eggs are present in the ovarioles as the adult molt and larva body mass is proportional to fecundity (i.e., 'capital breeders', (Honěk, 1993; Awmack & Leather, 2002)). As a consequence, host plant quality during larval growth and development is the key determinant of individuals fitness (Awmack & Leather, 2002). Although the relationship between plant quality and herbivore fitness may vary among species (Moreau et al., 2006; Awmack & Leather, 2002; Colasurdo et al., 2009), we speculate that herbivory by the first BTM larva generation reduces the fitness of the second BTM generation, and that this effect may be further strengthened where high population density increase intra-specific cross-generational competition (Tammaru & Haukioja, 1996).*

3. Minor comments:
4. I would remove "multivoltine" from the title. The cross-generational already gives the idea....

**[R] — Done**

5. Line 19: replace "proposed" by "offered".

**[R] — Done**

6. Line 23: unclear if this number of eggs is from the previous or the current brood.

**[R] — Changed**

7. Line 37: replace "their" by "its".

**[R] — Done**

8. Line 54: "deters" instead of "deter".

**[R] — Done**

9. Line 60: incidentally, previous herbivory can also lead to increased performance in subsequent infestations (e.g., Sarmento et al 2011 Ecol Lett, Godinho et al 2016 Oecologia). This is just a side comment, you don't need to include this in the paragraph...

**[R] — Thank you for these references.**

10. Lines 94-100. This paragraph is a bit confusing. First, I suggest placing the sentences on the biological details of the system (lines 96-98) elsewhere, maybe in the very beginning of the Material and Methods section. Second, it is not very clear to me when were the moths placed on the experimental trees. Is "the overwintering generation" the same as the "caterpillars collected in the wild"? If so, please be clear about this. Also, I guess that by "their" adults you mean "the adults emerging from those larvae"? (also commented by one of the reviewers).

**[R] — We restructured this paragraph, adding a new subsection ("*natural history*") and clarifying the description of first and second BTM generations (section "biological material").**

11. Line 104: I would state "plants" instead of "plant individuals".

**[R] — Done**

12. Line 111: you seem to use "chrysalis" and "chrysalids" interchangeably. If these terms refer to the same thing (I guess they do...) please choose one.

**[R] — Done**

13. Results: Did you count the number of egg clutches per plant with at least one clutch or per experimental plant in general? That is, could there be a difference in the number of plants with no egg clutches among treatments?

**[R] — We screened every single plant and counted the number of egg clutches on all plants. We added the information on the presence/absence of eggs to the '*results*' section.**

14. I would not discard the data concerning chrysalids so easily. It is indeed a pity that you cannot discriminate whether more larvae died in one treatment vs the other or if there was active migration, as you state. But in any case, this means that more individuals of this second 'generation' are eventually found on the previously clean plants, and this is an interesting result per se. I would at least discuss this a bit further in the Discussion.

**[R] — We do agree this is very unfortunate we have not been satisfyingly efficient in preventing larvae from moving among plants. Although we have been reluctant to present this data in first instance, we now reinjected them back into the manuscript, with words of caution in the discussion. Changes can be seen throughout the text in the "methods / analyses", "*results*" and "*discussion*" sections.**

15. Lines 180-183: Maybe rephrase as to use a more fluid text style. Ex: One possibility for female BTM not choosing among plants may be that...

**[R] — Done. Indeed, it reads smoother.**

16. Lines 182-190: I think the main argument against this hypothesis comes from your own data: larval weight differs among treatments. Assuming this is correlated with fitness, there are consequences for the moths of their mothers' choice.

**[R] — Yes! thank you for mentioning this. We have added this obvious argument.**

17. Line 193: I don't understand why laying 200 eggs corresponds necessarily to a bet-hedging strategy. Maybe rephrase?

**[R] —Done.**

18. Lines 203-204: This paragraph is about the possible absence of cues, not about the possible absence of fitness consequences, so this sentence is best placed in the previous paragraph.

**[R] — This paragraph, starting with "*Prior box tree defoliation by the spring generation of BTM larvae reduced the performance of the next generation*" is about herbivore growth. We dealt with possible absence of cues in the previous paragraph.**

19. Line 206: I would remove "trait-mediated" from this sentence because I am not convinced that the dichotomy between the two explanations rests on this. Instead, I think that the two possible explanations are past vs current competition. Also is there a possibility to obtain the density of larvae in the two treatments? That is, the number of larvae per intact lead?

**[R] — We removed "*trait-mediated*". Although we acknowledge that this would have been a powerful way to further address competition, we did not precisely count the number of larvae per plant or per shoot, mostly to avoid disturbance.**

20. Lines 213-215: does this mean that larval weight has no effect on fitness / population growth? Please clarify.

**[R] — Done**:

> *the BTM is thought to have broad tolerance to variability in host traits, as suggested by previous observations that BTM larva growth rate did not differ significantly among box-tree varieties (Leuthardt et al., 2013). It is unknown whether herbivory induced changes in host traits are of the same order of magnitude as trait variability among varieties. However, assuming variability among varieties is greater, this result goes against the view that reduced performance of larvae of the summer generation resulted from box tree response to prior herbivory*

21. Line 219: although I agree with one of the reviewers that the possibility that food shortage may lead to cannibalism is fascinating, I would remove this sentence unless you have hard data on which to base this statement. In particular, if this were to be true, you would need to explain (a) why you still find the same overall number of larvae alive between the two treatments and (b) whether it is expected that

this cannibalism does not compensate for food shortage in terms of larval weight. Overall, I think that this observation opens too many doors, so either it is solid or it better be left out of the Discussion.

**[R] — We deleted the mention to cannibalism, because this is true we do not have hard data ta back it up.**

22. Lines 223-225: These sentences fit best in the next paragraph.

**[R] — We completely modified the corresponding paragraph to account for the several comments on the results and discussion.**

23. Line 245: replace "in particular to plants" by "in particular to those".

**[R] — Done.**

24. Lines 359-361: please check formatting here.

**[R] — Done.**

---

Reviewed by Inês Fragata, 2020-09-08 23:55

In this manuscript the authors test whether female choice for oviposition impacts intraspecific competition across generations. In order to do this, the authors compare oviposition and larvae weight of box tree moths on box trees previously exposed to conspecific herbivores or un-attacked controls. They observe that previous herbivory does not affect where female choose to lay eggs, but it affects larvae weight. This suggests that there is a mismatch between female choice and larvae performance, which is against the preference-performance hypothesis. The question that the authors are trying to answer is very interesting and can help us to understand better how species avoid intra and interspecific competition, even across different generations. Unfortunately, the methodological problems with the chrysalids made it more difficult to fully explore the potential of the question and experimental design. I have some questions/suggestions that may allow to explore better the data set that you have here, and go a bit deeper into your questions.

**[R] — Thank you for comments.**

Questions/suggestions:

- If I understood correctly, you have the percentage of damage per plant that was done by the first infesting larvae. You could use this as a covariate in your choice experiment to see if damage was a better way of predicting female choice, in addition to your spatial correlation structure. I think it could also be interesting to use the initial percentage of herbivory on the analysis of the larvae weight.

**[R] — We have now added this information and re-ran models accordingly. We therefore made appropriate changes in the *methods* | *Experimental design* and *Results* sections (but did not list all changes here). The reason we did not consider herbivory data in first instance was that potted plants were initially used in a completely separate experiment, for another purpose. We could not match every pot tags between the two projects, leading to missing data. However, because we agree that testing BTM response to actual herbivory rather than to a qualitative attacked/non-attacked factor, we decided to bring original data back into the main text.**

- Besides the effect on choice of the egg laying females, herbivory could also affect plant quality and manifest in other life stages. In addition to looking at the weight of the caterpillars, since you have a measure of egg to caterpillar mortality, it would be interesting to see if egg to larvae mortality was higher in attacked or control trees.

**[R] — We agree that this would have been a great addition to the paper. However, we only counted and measured a subset of larvae and chrysalis (up to five), and therefore are not able to follow this advice**

- Regarding the data on the chrysalids, why do you assume that it was the number of chrysalids that was wrong, and not the number of eggs or larvae (i.e. you could miss some larvae/eggs)? Does this excess occur more in non-damaged trees? Also, does the number of extra chrysalids match the number of missing caterpillars from nearby trees/attacked trees? Because it would be interesting to see if there was more dispersal for pre-attacked trees than for control trees. It is quite striking to have half your controls and 1 one attacked plant where this happens, so I wonder if there is something biologically interesting underlying this pattern. However, if you are planning to not analyse the data or speculate on it, I think it would be better to remove the chrysalid part, as the usefulness of the information is unclear.

**[R] — Please see our response to Dr Magalhães, above***

- L148 - Did you release the moths in this region of the plots (between the 1 and 3)? because that could be a reason for the spatial structure to occur? alternatively did it had a source of light/heat or something alike? because it is rather strange that they clustered around that region.

**[R] — We released moths at the four corners of the experiment to reduce the risk of spatial aggregation (information now added to the manuscript). We have no data to support any explanation regarding the aggregation of eggs in one particular part of the experiment. This could actually be because of light (the part of the greenhouse received more sunlight in the afternoon) or because of fresher air arriving from the doors.**

- L152 – Why did you use these different spatial correlation structures? and what does it mean to have these different spatial correlation structures? This is important to explain what are you accounting for in the analysis.

**[R] — We simply followed textbook recommendations when there is no *a priori* hypothesis on the shape of potential patterns.**

> *we had no particular hypothesis regarding the shape of the spatial correlation structure. We therefore ran separate models with different spatial correlation structures (namely, exponential, Gaussian, spherical, linear and rational quadratic), and compared them based on their AIC (Zuur, 2009)*

- L163:166 – From table 2 you have 3 models (and not two) that have similar performance, and they are not significantly better or worse compared to the quadratic one, as you need at least a difference of 2 in the AIC, using the rule of thumb from Burnham & Anderson 2004

**[R] — Thank you for noticing, we corrected the text.**

- In the first section of the discussion, I think two hypotheses that you don't mention are that 1) the moths may need cues from other life stages, such as female conspecific oviposition or the chrysalids; 2) you let too much time pass and the cues related with the conspecific were not present anymore.

**[R] — Thank you for these suggestions. We adapted the manuscript accordingly. As for (1):**

> *or that female BTM were indifferent to them at the time we conducted the experiment.* [...] *It remains however possible that BTM adults use other cues to select their hosts such as the presence of conspecific eggs, larvae or chrysalis.*

**as for (2)**:

> *it is also possible that induced defense reactions were delayed in box trees, or that they were already relaxed when we released BTM moths three weeks after the herbivory treatment (Karban, 2011), which remains to be evaluated.*

Text comments

- L17:20 – This sentence is a bit weird, suggestion: "We tested this hypothesis in a choice experiment with box tree moth females (*Cydalima perspectalis* Walker, Lepidoptera: Crambidae). These females were exposed to box trees (*Buxus sempervirens* L., Buxaceae) previously defoliated by conspecific larvae earlier in the season."

**[R] — Changed.**

- L30 – remove the thus from "Insects may thus reduce"

**[R] — Done.**

- L32 – I think you mean assumes instead of supposes

**[R] — Changed.**

- L38 – I don't understand why you say "in particular" here, are those the only traits that will be important for females to detect that correlate with larval performance? for example presence of predators and competitors could be other factors that females may detect before ovipositing.

**[R] — We replaced *in particular* by *for instance.***

- L41 – "time-lagged consequences on the preference"

**[R] — Done**

- L49 – I would substitute a mix of, with both

**[R] — Done**

- L60 – I would replace the "later herbivores" by later arriving/appearing herbivores

**[R] — Done**

- L61 – I would replace the "late coming herbivores" by later arriving/appearing herbivores

**[R] — We preferred keeping this sentence unchanged to avoid repetition with the previous one.**

- L71:73 – This last sentence is not very clear. Maybe: "Thus, in order to quantify the effect of prior herbivory on subsequent herbivore performance, we need to assess how it affects both female choice and progeny performance in attacked and non-attacked hosts."

**[R] — Done**

- L99:100 – "Their adults. . . " whose adults? you mean that the larvae were used on the preference test and the adult stage on the performance test? maybe something like: "The adult stage of these larvae were used in. . . "

**[R] — we clarified this point:**

> _We initiated BTM larvae rearing with caterpillars collected in the wild in early spring 2019, corresponding to those that had overwintered**. We reared them at room temperature in 4320 $cm^3$ plastic boxes, and fed them_ ad libitum_, *with branches collected on box trees around the laboratory. We used the next generation larvae to induce herbivory on box tree plants (experimental treatment, see below) and the subsequent adults for the oviposition experiment.*

- L112 – feed on missing a space

**[R] — Done**

- L117 –Any specific reason for waiting the three weeks? is it the amount of time that they would take to lay eggs again?

**[R] — We have added this information to the revised version:**

> *In addition, at this time, plants in the herbivory treatment had been cleared of caterpillars for three weeks* ***(corresponding to the duration of the chrysalis stage)*** *during which they were watered every two to three days from above.*

- L132 – Why did you wait 24h for weighting them? and not measured them right away? Also was there a high variance in larvae weight?

**[R] — We have added this information to the revised version:**

> *We kept them in Petri dishes without food for 24h* ***to make larvae empty their gut*** *and weighted them to the closest 10 μg.*

**We present variability in larval weight in the *'Results'* section:**

> *The mean weight of BTM larvae varied from 6 to 54 mg (mean ± SD: 20 ± 9 mg).*

- L140 – 61 instead of 60 (or otherwise you have the number wrong above)

**[R] — Changed**

- L140:141 – maybe "and only 1 previously attacked plant" instead of "(and only in 1 previously attacked plant)"

**[R] — We rephrased this sentence.**

- L146 – x and y coordinates of what?

**[R] — Changed**

> *We ran a generalized least square model (GLS) testing the effect of potted tree location in the experimental design (through their x and y coordinates, Figure 2) on the number of clutches per plant (*log*-transformed) from which we explored the associated variogram using the functions gls and Variogram in the nlme library.*

- L166 – I would not call them competing models. Additionally, you should add whether they show similar results, since you cannot say which one is best.

**[R] — *x* and *y* coordinates referred to the design of the experiment (Figure 2). We referred to '*competing models*' after Burnham & Anderson textbook (2002). We now state that the results would have been the same regardless of the spatial correlation structure. However, because this information is not essential, we preferred not reporting the detailed model outputs. Interested or skeptical readers will have access to raw data and codes and will be able to simple uncomment the corresponding lines of codes.**

- Fig1 – I would put this figure as supplementary material.

**[R] — We agree that this figure is not essential, but on the other hand we value this kind of illustration showing what the experiment looked like, because the reality is sometimes substantially different from what a '*Methods*' section give to imagine.**

- Fig 2- I would like to know what are the x and y axis? meters? random unities?

**[R] — Pots were installed 40 cm apart. The numbers on axes were misleading. We removed them.**

---

Reviewed by Raul Costa-Pereira, 2020-09-06 15:59

Castagneyrpol et al. present results from a well-designed experiment aiming to test the detrimental effects between conspecific individuals that have never meet. They studied a peculiar system where consumers (box tree moth larvae) have limited mobility, relying on their mother's oviposition decision to occupy good food patches (i.e., host plants). Interestingly, food patches are dynamic, and consumers can reduce resource quality to next-generation conspecifics by triggering defensive responses in host plants. Therefore, even if conspecific individuals never coexisted on the same individual host plant, the legacy of past "tenants" can reverberate negatively on current and future ones. Motivated by this interesting conceptual basis, the authors set up a greenhouse experiment to test how past herbivory affects preference (i.e., selection of oviposition sites by mothers, quantified as the number of egg clutches) and performance (i.e., individual consumer growth, quantified as average larvae body mass). Surprisingly, mothers did not avoid laying eggs on plants previously consumed by larvae, but larvae feeding on these plants with the legacy of past conspecifics were smaller. These results are exciting because they shed new light on the mechanisms shaping temporal dynamics of antagonistic interactions both between (plant-herbivore) and within (herbivore-herbivore) species.

**[R] — Thank you for this very nice summary and positive appreciation!**

I enjoyed reading the manuscript and think it is well-written and the figures are well-presented. The experimental design is creative and statistical analyses are solid (I particularly appreciated how the authors accounted for the underlying spatial structure of their experiment in the models). Below I describe a few major points that came up while reading the manuscript, as well as some minor points that I believe can be helpful. Please let me know if you have any questions, I am happy to clarify.

**[R] — Thank you for your valuable and clear comments.**

First, I feel that the conceptual framing of the manuscript is fascinating and could be contextualized and motivated more broadly in the introduction. Indeed, multivoltine insect herbivores are a great example of how individuals can affect conspecifics they have never met. Still, similar types of time-lagged interactions between conspecifics occur across diverse taxa (e.g., squirrels [Fisher et al. 2019 Ecology Letters], frogs [Pfennig & Pfennig 2020 Copeia]) and via different mechanisms (e.g., extend phenotypes, ecosystem engineers). Therefore, although the current structure of the introduction works well, I think that opening the manuscript with a more general view of ecological interactions among individuals separated in time would call the attention of a wider and more diverse readership. This approach could also help to reduce some overlap in ideas across the 1st and 2nd paragraphs of the introduction.

**[R] — We really appreciated this suggestion. We added a couple of opening sentences to broaden the scope of the paper**

> *Biotic interactions are strong factors affecting the fitness of interacting individuals, even interactions are delayed in time and do not imply direct contact among individuals. Examples of such interactions can be found in both plants through plant-soil feedbacks (Putten et al., 2016) and in animals (Fisher et al., 2019; Pfennig & Pfennig, 2020)*

Hypotheses and respective predictions could be more thoroughly presented to readers. The last paragraph of the introduction is concise and nicely describes the general hypotheses of the study (lines 84-85). However, readers will only find out how the authors investigated their hypotheses in the methods (e.g., lines 123-135), which creates a certain gap in the narrative flow. Thus, the authors could include their respective predictions as well at the end of the introduction (including 'operational variables' - e.g., We expect that plants that previously hosted larvae should [i] have fewer eggs and [ii] host smaller larvae). Moreover, as the experimental design allows inferring the contribution of purely spatial effects on oviposition patterns, the authors could at least mention this at the end of the introduction. By the way, I think that measuring and accounting for spatial structure in oviposition patterns is an exciting novelty of the manuscript. Thus, maybe the biological causes and implications of this spatial non-independence in mothers' oviposition choices could be more explored in the discussion.

**[R] — We have modified the end of the introduction in order to introduce "*operational variables*" as suggested. As for the discussion on spatial analyses, we agree that our results could pave the way for further investigation. However, the experiment was not designed to explore such spatial effects. We only aimed at controlling possible bias in the design. Therefore, we preferred not putting to much emphasis on this issue as it would have been very speculative**.

Finally, I have a quick suggestion about a potential additional dimension of performance that could be considered. To quantify the effects of past herbivory on offspring performance, the authors compared the average body mass of larvae across treatments. I fully agree this is a key aspect of *per capita* offspring performance, and results are exciting in this regard. In light of the natural history of the system, as the authors also measured the number of L3 larvae per plant (lines 130-131), I wonder whether the conversion rate egg → L3 larva (e.g., number of eggs/number of L3 larvae) could not be used as an additional metric of performance. Is there evidence in the literature that past herbivory can affect egg eclosion rates and/or early larval development (L1 → L2 → L3)? This alternative metric would capture a different facet of offspring performance not necessarily correlated with mean larvae weight (e.g., larvae mortality associated with lower foraging rates and/or increased toxins).

**[R] — This would have been a great addition to the paper indeed. Unfortunately, we did not have such an information at hand for we counted *up to* five larvae per plant. The phrasing of the original version was ambiguous in this respect. We modified it accordingly.**

Minor comments:

- Lines 1-2. The title is solid and general, but I am not sure if all readers will be familiar with the concept of 'multivoltine' (I guess it depends on the target journal).

**[R] — We deleted the reference to *multivoltine* species**.

- Lines 39-40. Given the idea of the last sentence in this paragraph, it would be good to emphasize here that "competing herbivores" refers to different species of herbivores (i.e., interspecific competition). I would say the same about Line 42.
- Lines 46-50. I feel that most of these ideas were already presented to readers in the previous paragraph. The argument presented in Line 50 sounds like an exciting way to begin this paragraph.

**[R] — We deeply modified the introduction, please see our response to Dr Magalhães' comments.**

- Line 50. I follow the meaning of 'passage of competitors' but it's possible that some readers may find it a bit confusing. One potential alternative (maybe not that accurate) would be something like 'the legacy' of past herbivores on host plants.

**[R] — The corresponding sentence was deleted**.

- Line 51. Maybe the authors could provide a brief view of what 'direct' detection means in this context, e.g. "…mated females can directly detect (e.g., via visual or olfactory cues) the present…".

**[R] — We added "themselves" to make it clear that the female can detect herbivores, or herbivory-induced changes in plant traits**.

- Line 61. Adding a 'triggering' or 'stimulating' before "defenses that generally" could make this idea clearer to readers.

**[R] — We rephrased this sentence**.

- Line 60. I am not an expert in plant-herbivore systems, but a first intuitive, simple mechanism seems to be the reduction of food biomass by previous consumers. Does it make sense?

**[R] — Yes it does! We now mention interference competition and resource depletion**.

- Lines 65-66. This is very interesting, and the following example illustrates well this mismatch between selection by mothers vs. impacts on offspring. However, I feel a follow-up conceptual sentence would help readers to crystallize this idea by clarifying that not necessarily the effects on preference and performance are congruent.

**[R] — We restructured the introduction to improve the reading**.

- Lines 74-75. Considering the broad readership of ecologists and evolutionary biologists this manuscript has the potential to reach, I recommend the authors to define the concept of 'multivoltine'. Not all readers may be familiar with it.

**[R] — Done**.

- Line 75. Tiny detail: remove the italic from spp.

**[R]** — **Done**.

- Lines 84-86. This is a matter of writing style, but one possibility here is to 'change the pace' of this key sentence to emphasize the potential effects on both preference and performance encapsulated by this hypothesis. A simple way to do this would be: "... early herbivory would (i) reduce oviposition..., and also (ii) reduce the performance ...

**[R]** — **Done**.

- Lines 85-86. As the last sentence of the introduction is often one of the most 'visited' by readers, I feel this one could deliver a 'self-standing', stronger message. For instance, instead of 'By addressing the above', one alternative could be 'By addressing the effects of previous herbivory by conspecifics on both preference and performance of subsequent... "

**[R]** — **Done**.

- Line 94. I think this initial sentence could be moved down in this paragraph.

**[R]** — **The sentence was moved to the "natural history" new paragraph.**

- Line 107. It is clear from the previous sentence, but it would be helpful to clarify that this plant-level herbivory metric represents the mean frequency of attacked leaves/branch.

**[R]** — **Done**:

> _ In order to confirm that the addition of BTM larvae caused herbivory, we visually estimated BTM herbivory as the percentage of leaves consumed by BTM larvae, looking at every branch on every plant. We then averaged herbivory at the plant level. In 8 plants, herbivory data was missing and was imputed as the average of herbivory measured in other plants. In the herbivory treatment, the percentage of leaf area consumed by BTM larvae ranged from 2.2 to 17.2% and was on average 9.1%._

- Figure 1. These photos are great for illustrating the experimental design and study system! I would just suggest adding more details in the legend.

**[R]** — **Done**.

- Line 113. feed on.

**[R]** — **Done**.

- Lines 115-116. I wonder if this difference of three weeks between caterpillars being removed (from the herbivory treatment) and moths oviposition reflect the phenology of this species in natural ecosystems. In other words, in light of the biology of BTM, a given box tree in nature could experience two separate groups/generations of caterpillars within three weeks?

**[R]** — **Yes, the duration of the pupal stage in the wild is long enough to have two separate generations**.

- Line 118. Just to make it more straightforward: "... washed out from leaves".

**[R]** — **We deleted this sentence**.

- Line 120-121. This is an important point that could be briefly mentioned in the last paragraph of the introduction, e.g., "our experimental design allows us to quantify... of plant-mediated".

**[R]** — **Done**.

- Lines 126-128. I think that first presenting 'why' (i.e., "to prevent larvae from moving from one potted plant to another") and then 'how' (i.e., "we installed box trees in plastic saucers and interspaced plants and filled saucers with a few centimeters of water").

**[R]** — **Done**.

- Line 131. All plants had at least five larvae?

**[R]** — **Done: "only 6% of plants hosted less than five larvae".**

- Line 133. It would be helpful to emphasize that this value represents the average weight of larvae from one plant individual.

**[R]** — **Done**.

- Line 142. Thanks for such a careful explanation; I appreciate your transparency!

**[R]** — **Thanks**.

- Line 150. Could you please describe in more detail the structure of these models? (i.e., define response variable and predictors).

**[R]** — **Done**.

- Line 165. With other three models ($\Delta i < 2$ - Models 2, 3, and 4), right?

**[R]** — **Yes, corresponding information is now available in Table 1**.

- Line 177. How about finishing this sentence after 'season' and then start a new sentence with 'This time-lagged'?

**[R]** — **Done**.

- Line 182-183. Maybe the fact that larval frass was washed out could be a potential explanation?

**[R]** — **We added the following sentence:**

> *However, we cannot exclude that some cues were mediated by larva frass, which was watched out from leaves when we watered plants.*

- Lines 201-203. This an interesting explanation! I wonder if host plants in the native range of BTM have even higher toxic alkaloids than box trees.

**[R]** — **This is an interesting question that would be worth digging further. We are not aware of dedicated studies**.

- Lines 206-207. I do not follow this idea, could you please clarify? I feel that 'reduced performance of individuals... have been trait-mediated' requires some further explanation.

**[R]** — **"Trait mediated" was misleading. We deleted these two words and believe the sentence reads better now**.

- Line 219-220. The fact that food limitation can trigger cannibalism in this system is fascinating!

**[R]** — **yes, we have been quite surprised to observe this, but did not try to investigate it further. However,in the absence of back-up data, we eventually deleted reference to this possible phenomenon**.

- Line 221. To avoid repetition, I suggest replacing the first 'Herbivore' by 'Consumer'.

**[R]** — **This sentence was deleted**.

- Line 228. Would it be 'with' or 'within'?

**[R]** — **Changed to "within"**.

- Line 237. Perhaps 'negative interactions... generations' could communicate more clearly the results.

**[R]** — **Done**.

- Line 241. Because this idea expands to the next sentence, it would be good to mention their main enemies (e.g., parasitoids, predators).

**[R]** — **Done**.

- Line 244. It seems that a verb is missing in this sentence, 'causing more damage' is one option.

[**R**] — **Done**.

- Line 247. be investigated

[**R**] — **Done**.

- Line 248. dedicated

[**R**] — **Done**.

I hope the authors find these comments helpful. Best wishes, Raul.

[**R**] — **We did, thanks!**

#### 3.2 Second round

Dear authors,

I think you made a great job at replying to the concerns of the referees and opted not to send them the manuscript again. I do have some very minor comments still that I think should be addressed in the version of this manuscript that will then be accepted. Congratulations!

[**R**] — **Thank you for your very thorough review and positive comments on the revised version of our manuscript.**

- Lines 52-53: I would start general, stating that previous herbivory can affect the performance of subsequent herbivores. Then, when later on you refer to the effect via plant defences you mention the possibility of actually facilitating future herbivory. I think this part is important because it adds a bit of unexpected outcome to your story. Otherwise, the prediction of previous herbivory being detrimental is a bit too straightforward.

[**R**] — **We reorganized the corresponding paragraph accordingly:**

> *Previous herbivory generally reduces the performance of later arriving herbivores on the same plant through different processes. First, the initial consumption of plant biomass can deplete the resource available to forthcoming herbivores, therefore leading to exploitative competition between first and subsequent herbivores (Kaplan & Denno, 2007). Second, initial herbivory triggers a hormonal response that results in the induction and production of anti-herbivore defenses as well as in resource reallocation in plant tissues (Hilker & Fatouros, 2015; Abdala-Roberts et al., 2019; Marchand & McNeil, 2004; Blenn et al., 2012; Fatouros et al., 2012), which generally reduces plant quality and thereby the performance of late coming herbivores (Agrawal, 1999; Abdala-Roberts et al., 2019; Wratten et al., 1988). Such an affect has long been documented in interspecific interactions (Kaplan & Denno, 2007; Moreira et al., 2018), but also in intraspecific interactions. For instance, prior damage by the western tent caterpillar* Malacosoma californicum *Packard (Lepidoptera: Lasiocampidae) induces the regrowth of tougher leaves acting as physical defenses and reducing the fitness of the next tent caterpillars generation (Barnes & Murphy, 2018). Although less common, the opposite phenomenon whereby initial herbivory faciliates damage by subsequent herbivores has also been reported (Sarmento et al., 2011; Godinho et al., 2016; Moreira et al., 2018).*

- Line 58: a hormonal

[**R**] — **Done**

- Line 61: effect

[**R**] — **Done**

- Line 62: space between interactions and bracket

**[R] — Done**

- Line 66: previous herbivory can also affect the oviposition preference of herbivores that arrive later.

**[R] — We rephrased the original sentence.**

- Line 67: can discriminate between.

**[R] — Changed.**

- Lines 77-85: incidentally, we have data showing a match between preference and performance in absence of competitors, but not in their presence (Godinho et al. "The distribution of herbivores between leaves matches their performance only in the absence of competitors" Ecol Evol 2020). This is just a note, I think it's an interesting result that could be discussed in your paper, but you don't need to do it…

**[R] — This is a nice paper which we refer to in the revised version.**

- Lines 125-126: I would say that it is more parsimonious to simply remove those plants from the analyses. Can't you do that?

**[R] — We followed this recommendation. Ignoring these individuals slightly changed the results in terms of significance. Specifically, this new analysis revealed that the negative effect of prior herbivory increased with increasing the amount of leaf area removed by BTM larvae earlier in the season. We modified the results section accordingly and added one sentence in the discussion.**

> **Methods**: *Herbivory data were missing in 8 plants. We removed these plants from the analysis testing the effect of prior herbivory as a continuous variable on BTM preference and performance.*
>
> **Results**: *The effects of herbivory treatment and number of egg clutches on mean chrysalis weight were very comparable to those observed for BTM larvae: BTM chrysalis weight was lower on box trees that had been previously defoliated (**Table 2, Figure 3C**), and this effect strengthened with an increasing amount of herbivory.*
>
> **Discussion**: *This explanation is further supported by the fact chrysalis weight was more reduced in plants that were more defoliated by the spring generation of BTM larvae.*

- In lines 146 and 148, both sentences start with 'In order to', you can easily replace one of them by 'To'.

**[R] — Changed.**

- Figure 1: I actually agree with reviewer 2 that this fits better supplementary material, but you can also leave it, it's a matter of taste. However, the legend should be a bit more serious…

**[R] — We kept Figure one in the main text but rephrased the caption.**

- Line 161: were run.

**[R] — Changed.**

- Line 164: remove one of the brackets.

**[R] — Done.**

- Lines 191-192: this sentence was not in the original version of the manuscript, and I have difficulties in reconciling it with the previous sentence. If egg clutches were found in more than 90% of the plants overall, how can they, per treatment, be found on circa 40% of them?

**[R] — The percentage of plants with eggs was calculated over the total number of plants, and not the number of plants per treatments, which was indeed misleading. We clarified this issue in the revised version:**

> *We counted eggs in 93.4% of plants in the control (non attacked) groups, and in 100% of plants in the herbivory treatment.*

- Line 192. I don't understand why this sentence starts with 'however' nor why you are stressing 'at individual plant level'. Isn't this just the variance around the average numbers presented in the previous sentence? Maybe I am missing something important here...

**[R] — "However" was indeed not necessary, we deleted it.**

- Line 198: for the sake of consistency.

**[R] — Changed.**

- Line 201: I would say "Herbivory had no effect" because in the second analysis it is not the treatment per se that you are analysing.

**[R] — Changed accordingly.**

- Line 209: I think that what you mean is that 'larval weight was not significantly affected by the interaction ... ". Right? Please state this explicitly, I got a bit confused.

**[R] — Changed accordingly:**

> *Larval weight was not significantly affected by the interaction between the herbivory treatment and the number of egg clutches, indicating that intraspecific competition was independent of prior herbivory (**Table 2**).*

- Lines 220, 221: same here, I would merge the two sentences: There was a significant, negative relationship between the number of egg clutches on a box tree and the subsequent chrysalis weight, which was not significantly affected by the interaction between the herbivory treatment and the number of egg clutches (Table 2, Figure 3C).

**[R] — We rephrased the initial sentence accordingly.**

- Lines 227-230: please rephrase this sentence to clarify that you are proposing two explanations, not just one.

**[R] — Done:**

> *Prior herbivory had no effect BTM oviposition choice. Two possible mechanisms can explain this observation: prior herbivory may have had no effect on box tree characteristics, or female BTM may have been indifferent to them at the time we conducted the experiment.*

- Line 231: later, not latter.

**[R] — Corrected**

- Lines 240-242: why? Can larvae move among trees? If not, I don't think this is a proper explanation.

**[R] — This paragraph seeks to explain why prior herbivory had no effect on oviposition choice. Female BTM searching for oviposition sites were free to move in the greenhouse.**

- Lines 242-243: so what? This may be interesting, but you need to spell out your reasoning here. Do you mean that it may be more important to accumulate those alkaloids for the future reproductive success of those larvae than to be of a particular weight? This may be true, but still, they are not facing the choice of no alkaloids and big vs alkaloids and small, right? Or is there evidence that pre-attacked plants have more alkaloids than clean plants? This whole issue needs to be further developed or excluded...

**[R] — We explained this idea in more details:**

> *Leuthard* et al. *(2013) showed that BTM larvae are able to store or metabolise highly toxic alkaloid present in box tree leaves. Even if prior herbivory induced the production of chemical defenses, it is possible they this did not exert any particular pressure upon females for choosing undefended leaves or plants on which to oviposit, as their offspring would have been able to cope with it.*

- Line 248: add a coma after hosts.

[R] — **Done.**

- Line 263: I would remove the 'however'.

[R] — **Done.**

- Line 268: the 'interference competition' comes a bit out of the blue here. Is there any evidence for this in your system? And for the possibility of it being stronger in defoliated plants?

[R] — **The reference to "interference competition" was a mistake. We replaced it by "exploitative competition"**

- Lines 270-271: I would write this statement in the same style as the others, ie: The number of egg clutches laid by BTM female moths correlated negatively with subsequent growth of BTM larvae.

[R] — **Changed accordingly.**

- Line 280: was greater than.

[R] — **Changed.**

- Line 282: I would state 'this suggests' instead of 'indicates'.

[R] — **Changed.**

- Lines 285-287: I would remove this because you already state this in the previous paragraph.

[R] — **We removed this sentence.**

- Line 296: I would replace where by when.

[R] — **Changed.**

- Lines 297-298: I would state instead: These cross-generational effects may thus lead to an important role of density dependence population growth.

[R] — **Changed.**

- Line 307: add a comma after hosts.

[R] — **Done.**

- I find the conclusion still a bit too much attached to the system. I was wondering whether this can be linked to the Ghost of competition past. I will think about it a bit more and maybe write something on it in my recommendation, but maybe you can tell me what you think about this link before that...

[R] — **We completely re-wrote the discussion to make it more general. We did not phrase it in terms of "the ghost of competition past" but will love reading a recommendation about it!**

> *Insect herbivory induces changes in the amount and quality of plant resources, which are responsible for interspecific interactions among herbivores, even in herbivores that are separated in space or time (Poelman et al., 2008; Stam et al., 2014). Our experiment provides evidence that insect herbivory also influences the performance of conspecific herbivores through cross-generational competition, which may ultimately control the overall amount of damage that multivoltine herbivore species can cause to plants. Cross-generational competition may increase development time of individuals of the next generation, thereby increasing their vulnerability to natural enemies (the* slow-growth-high-mortality hypothesis*; Coley et al., 2006; Benrey & Denno, 1997; Uesugi, 2015). If this is the case, on the one hand stronger top-down control can be exerted on herbivores feeding on previously attacked hosts, which could reduce the overall amount of damage to the host plant. On the other hand, if herbivores take longer to develop, they may cause more damage to plants, in particular to those with poor nutritional quality, due to compensatory feeding (Simpson & Simpson, 1990; Milanovic et al., 2014). Our results highlight the overlooked ecological importance of time-lagged intraspecific competition (Barnes & Murphy, 2018). In the face of global warming, which*

*shortens the generation time of many insect herbivores and thus increases voltinism (Jactel et al., 2019), it is particularly necessary to elucidate the consequences of cross-generational interactions on the population dynamics of multivoltine herbivore species.*
