## Supplementary figures and images for "Host-mediated, cross-generational intraspecific competition in a herbivore species" (from another version of this preprint: version 4, 2021-02-11)

### Fig2-1.pdf

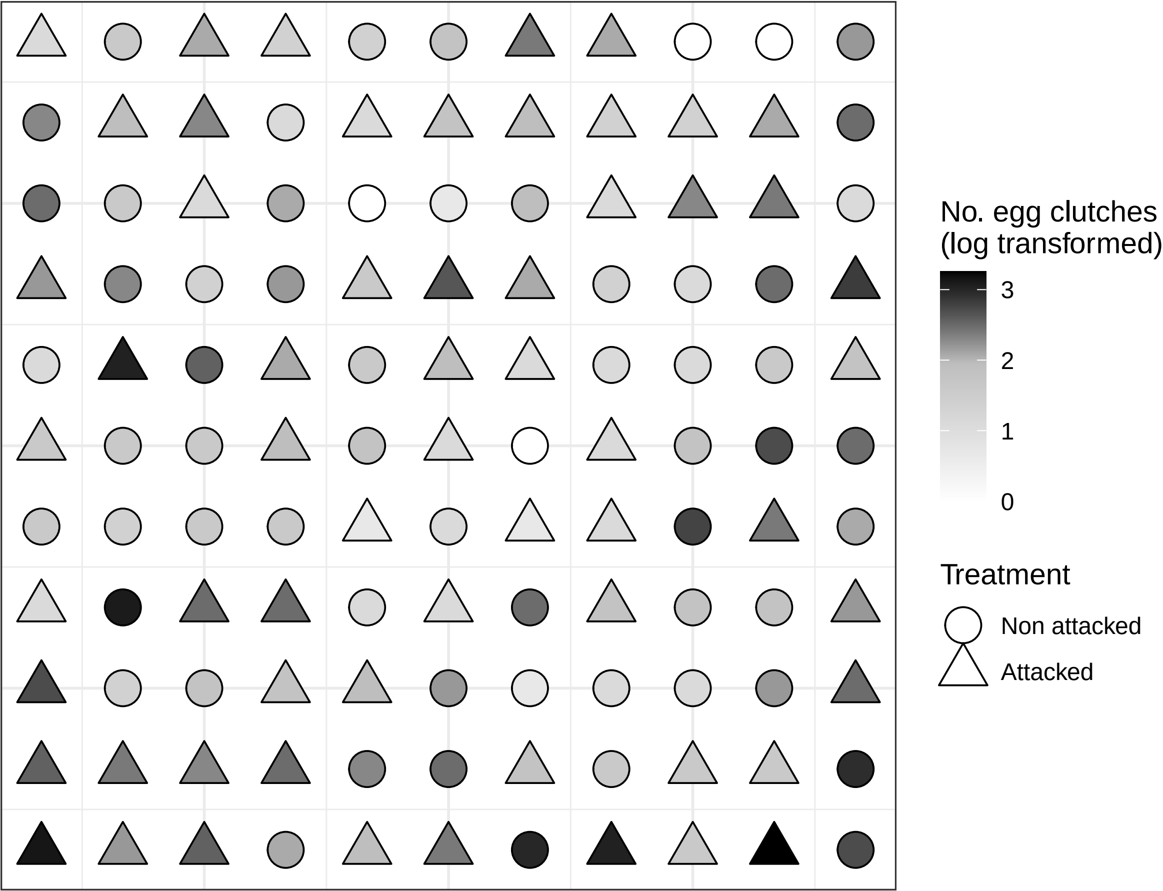

### Fig2-1.png

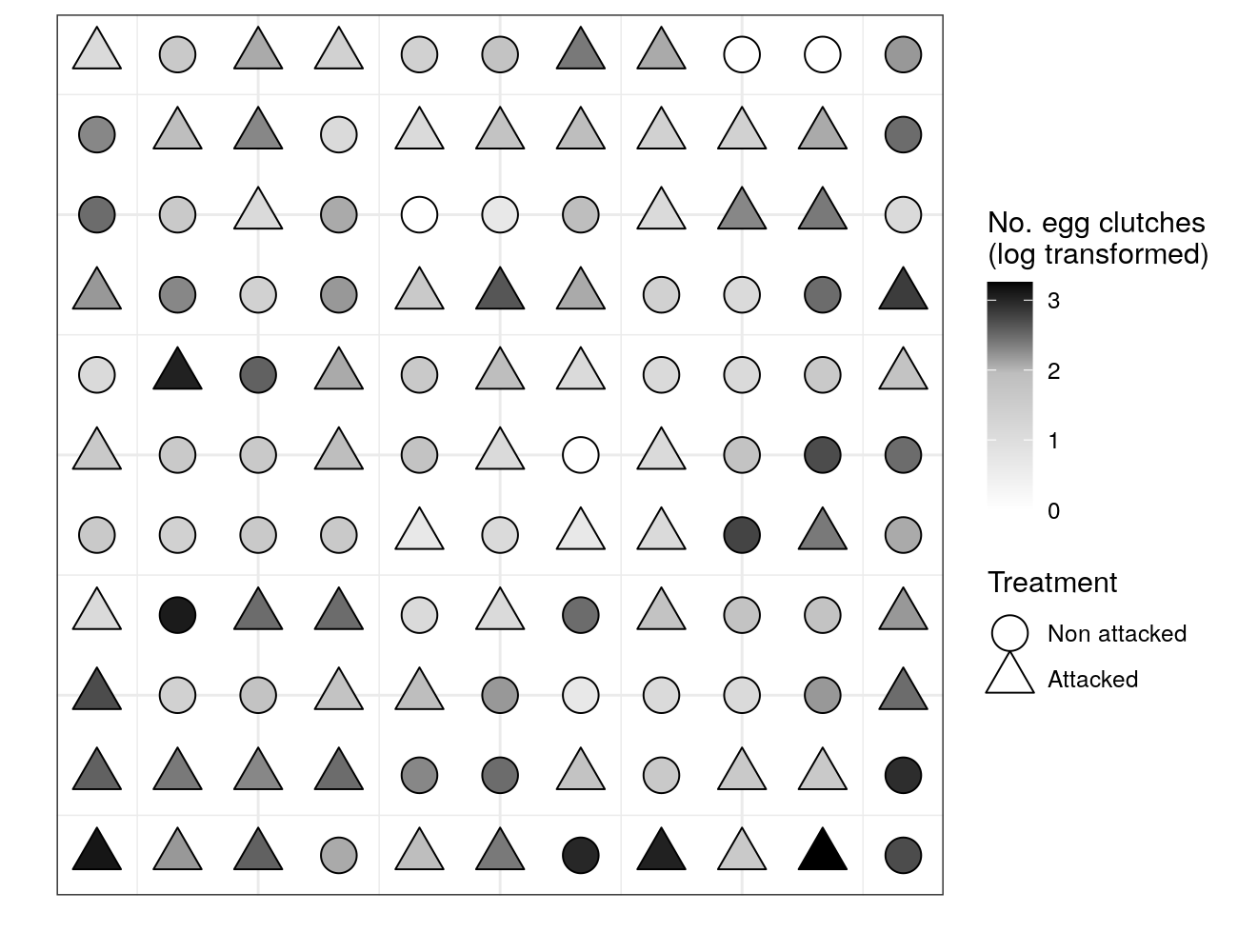

### Fig3-1.pdf

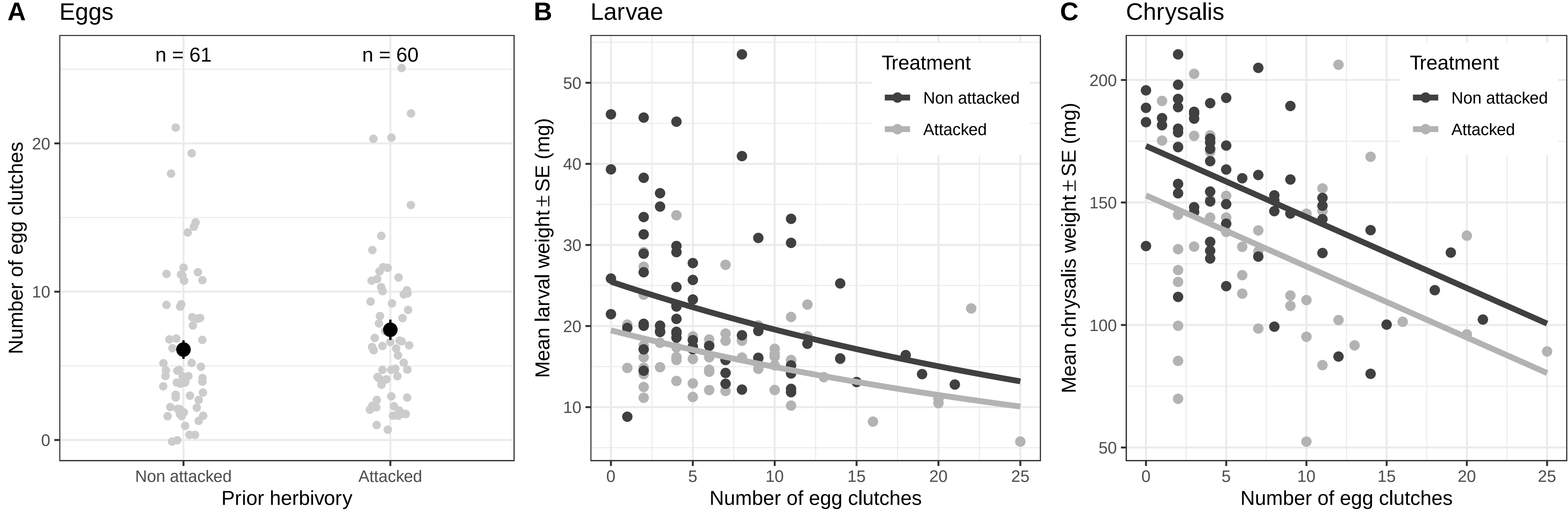

### Fig3-1.png

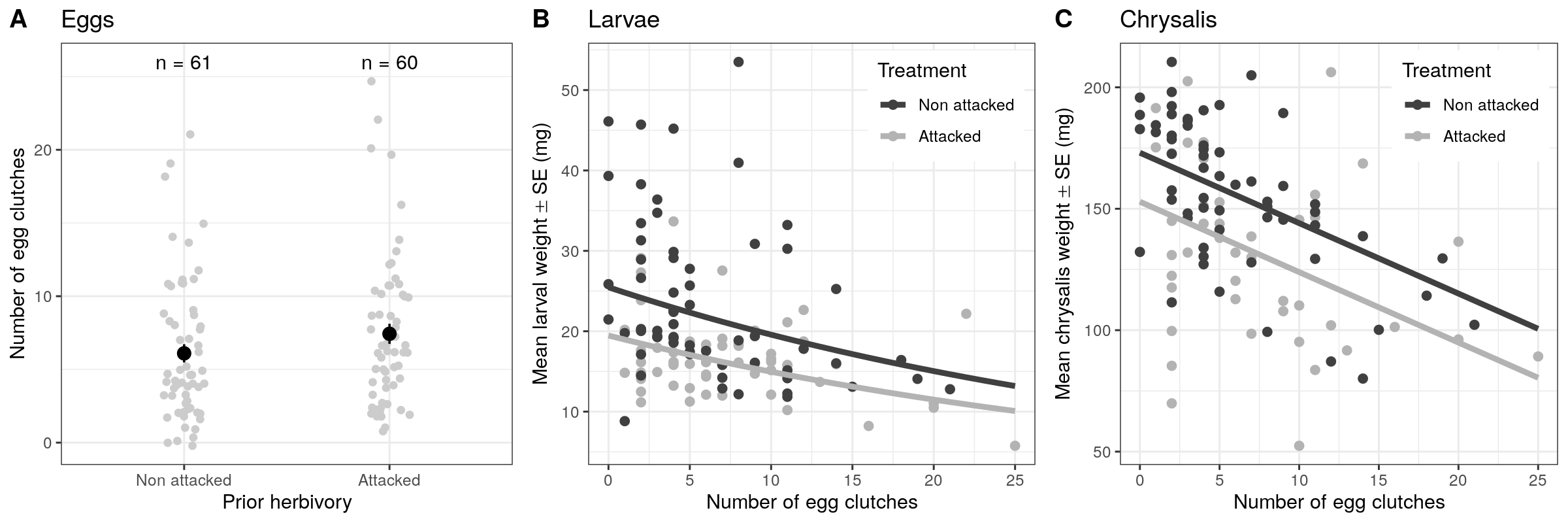

### FigS1-1.pdf

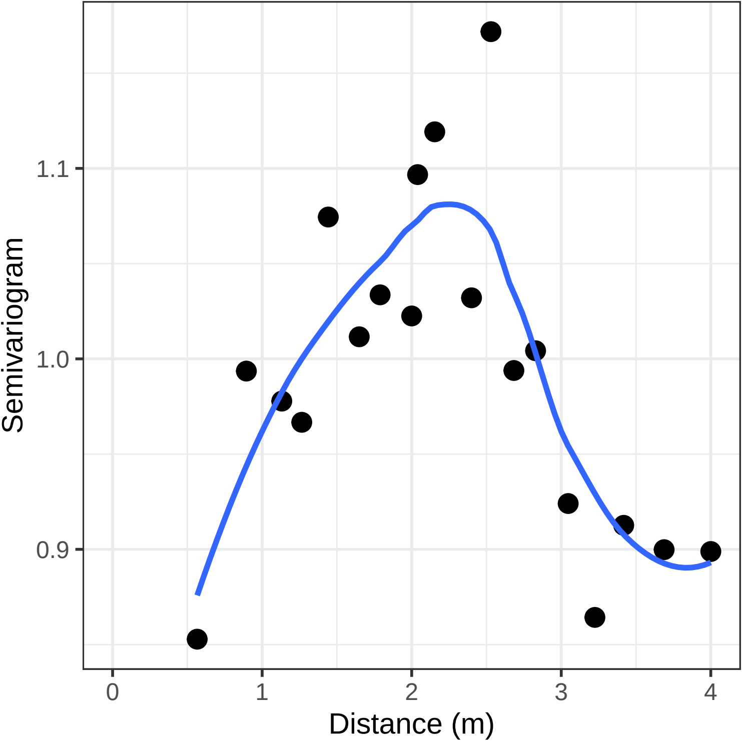

### FigS1-1.png

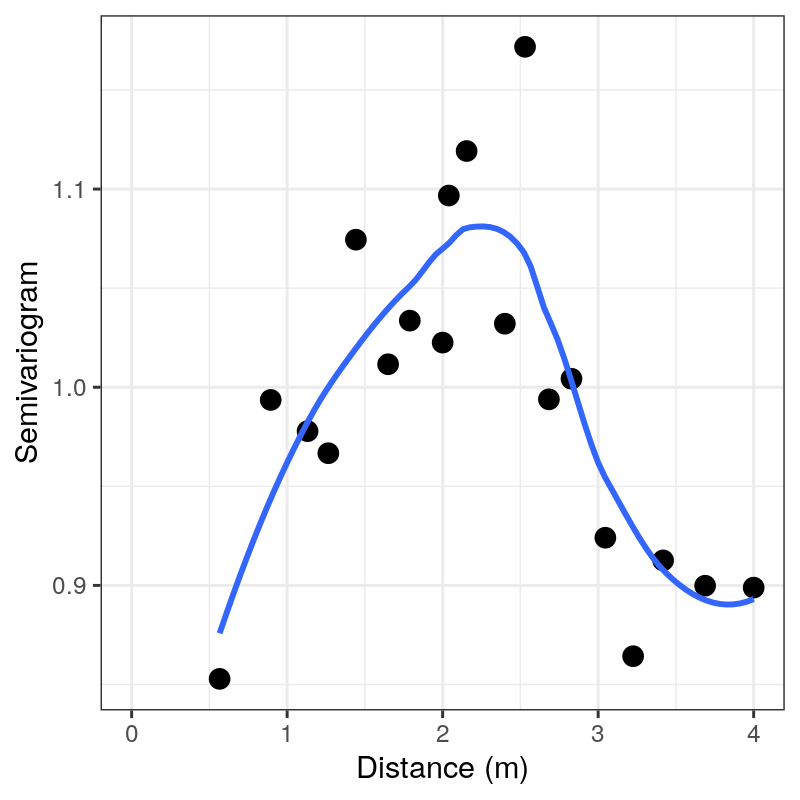

### Figure 1.png

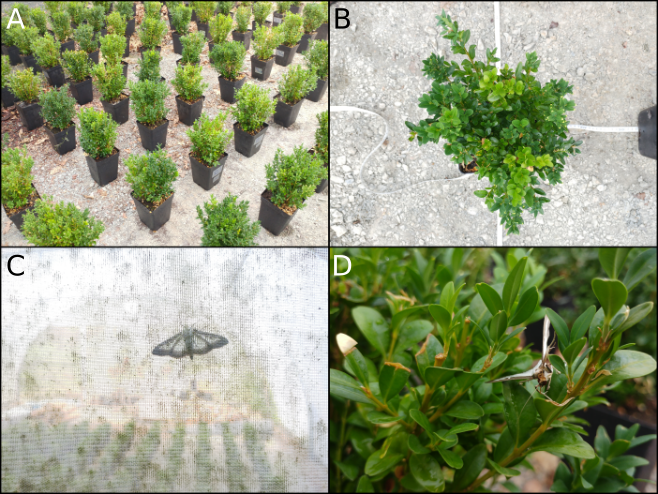
